# TM6SF2 is a cholesterol-bound homomultimeric ER membrane protein that binds ApoB and is required for bulk lipidation of VLDL

**DOI:** 10.64898/2026.08.17.745233

**Authors:** Sen Hong, Jin Wang, Matthew A. Mitsche, Jonathan C. Cohen, Xiaochun Li, Helen H. Hobbs

**Author notes:** Correspondence should be addressed to X.L. or H.H.H. or.

## Abstract

A missense variant in TM6SF2 (transmembrane 6 superfamily member 2, TM6SF2^E167K^) is a major risk factor for steatotic liver disease^1^, while protecting against coronary artery disease^2^. TM6SF2 is a polytopic resident protein of the smooth endoplasmic reticulum (ER) and ER-Golgi intermediate compartment that promotes lipidation of hepatic ApoB-containing lipoproteins before secretion into the circulation. Here, we used cryo-electron microscopy (cryo-EM) to determine the structures of TM6SF2 and TM6SF2^E167K^. TM6SF2 comprises 10 transmembrane helices that bind a single cholesterol molecule within a transmembrane cavity. The protein assembles into homodimers and homotetramers that interact with ApoB. Structural and biochemical analyses show that the E167K substitution reduces cholesterol binding and ApoB interaction without disrupting overall protein structure. Expression of wild-type, but not mutant, TM6SF2 restores hepatic triglyceride secretion in TM6SF2-deficient hepatocytes. Together, these findings establish the structural framework for the bulk lipidation step in hepatic lipoprotein biogenesis, the principal pathway for hepatic triglyceride and cholesterol export into the circulation.

## Introduction

Steatotic liver disease (SLD) is a prevalent metabolic disorder characterized by accumulation of triglycerides (TGs) within cytoplasmic lipid droplets (LDs) of hepatocytes. The disease develops when lipid influx from diet and peripheral tissues plus *de novo* lipogenesis exceeds lipid disposal through mitochondrial β−oxidation or export of TG as very low-density lipoprotein (VLDL)^3,4^. The global burden of SLD is substantial, affecting approximately 8%-11% of children^5^, up to 24% of adolescents^6^, and ∼38% of adults.^7^ Typically, the disease first presents with hepatic steatosis in individuals who are obese, have insulin resistant, or consume excess alcohol. In some affected individuals, steatosis is accompanied by inflammation and fibrosis,^3,4,8^ which can culminate in cirrhosis and/or hepatocellular carcinoma^3,9^. In epidemiological studies, SLD is strongly associated with coronary artery disease (CAD), although whether this relationship is causal remains unresolved.

Previously, we identified the two strongest genetic risk factors for SLD: PNPLA3^I148M^ ^10^and TM6SF2^E167K 1^. Homozygotes for either risk allele have approximately two-fold higher hepatic TG content than homozygotes for the corresponding nonrisk alleles. Both variants confer a similar risk of metabolic dysfunction-associated and alcohol-associated SLD.^4^ Despite these similarities, they differ in a key respect: PNPLA3^I148M^ is not associated with an increased risk of CAD, whereas TM6SF2^E167K^ is associated with protection from CAD^2^, likely due to its association with lower plasma LDL-cholesterol (C) levels.^2^

TM6SF2 is highly expressed in enterocytes and hepatocytes, both of which synthesize ApoB-containing lipoproteins. The protein resides in the smooth endoplasmic reticulum (ER) and the ER-Golgi intermediate compartment (ERGIC) of hepatocytes where it is required for bulk lipidation of newly synthesized VLDL^11–14^. *In silico* structural modeling predicts that TM6SF2 contains 10 transmembrane (TM) helices and two short, intralumenal α-helices (α1 and α2). The TM helices are tandemly arrayed, each harboring an EXPERA (EXPanded EBP superfamily) fold that resembles the sterol isomerase EBP (emopamil-binding protein)^15^.

To define the role of TM6SF2 in the synthesis of ApoB-containing lipoproteins, we determined the cryo-EM structures of wildtype (WT) mouse TM6SF2 (mTM6SF2^WT^) and the E167K variant (mTM6SF2^E167K^) at 3.6 Å and 3.58 Å resolution, respectively. Each protomer of the WT protein contains 10 TM helices and a cavity that accommodates cholesterol. The protein assembles into higher order homodimers and homotetramers. The E167K substitution compromises cholesterol binding to TM6SF2 and reduces its association with ApoB-100, the principal apolipoprotein of VLDL. Consistent with these structural and biochemical defects, TM6SF2^E167K^exhibits impaired activity in promoting VLDL lipidation when expressed in cultured hepatocytes lacking TM6SF2. Together, these results identify TM6SF2 as a cholesterol-dependent ER protein that is required for efficient VLDL-TG secretion.

## Results

### Structure of tetrameric TM6SF2

Full-length mouse TM6SF2 (mTM6SF2) with an N-terminal FLAG tag (Fig. 1a) was expressed in human embryonic kidney (HEK) 293S GnTI^−^ cells, and purified by size-exclusion chromatography (SEC) with glyco-diosgenin (GDN) (Fig. 1b, left). Two major peaks were observed, corresponding to the dimeric and tetrameric species, consistent with results obtained by Blue Native-PAGE (monomer M.W. 42 kDa) (Fig. 1b, right). Cryo-EM analysis further indicated that mTM6SF2 exists as a mixture of dimers and tetramers (Fig. 1b, left).

**Fig. 1.**
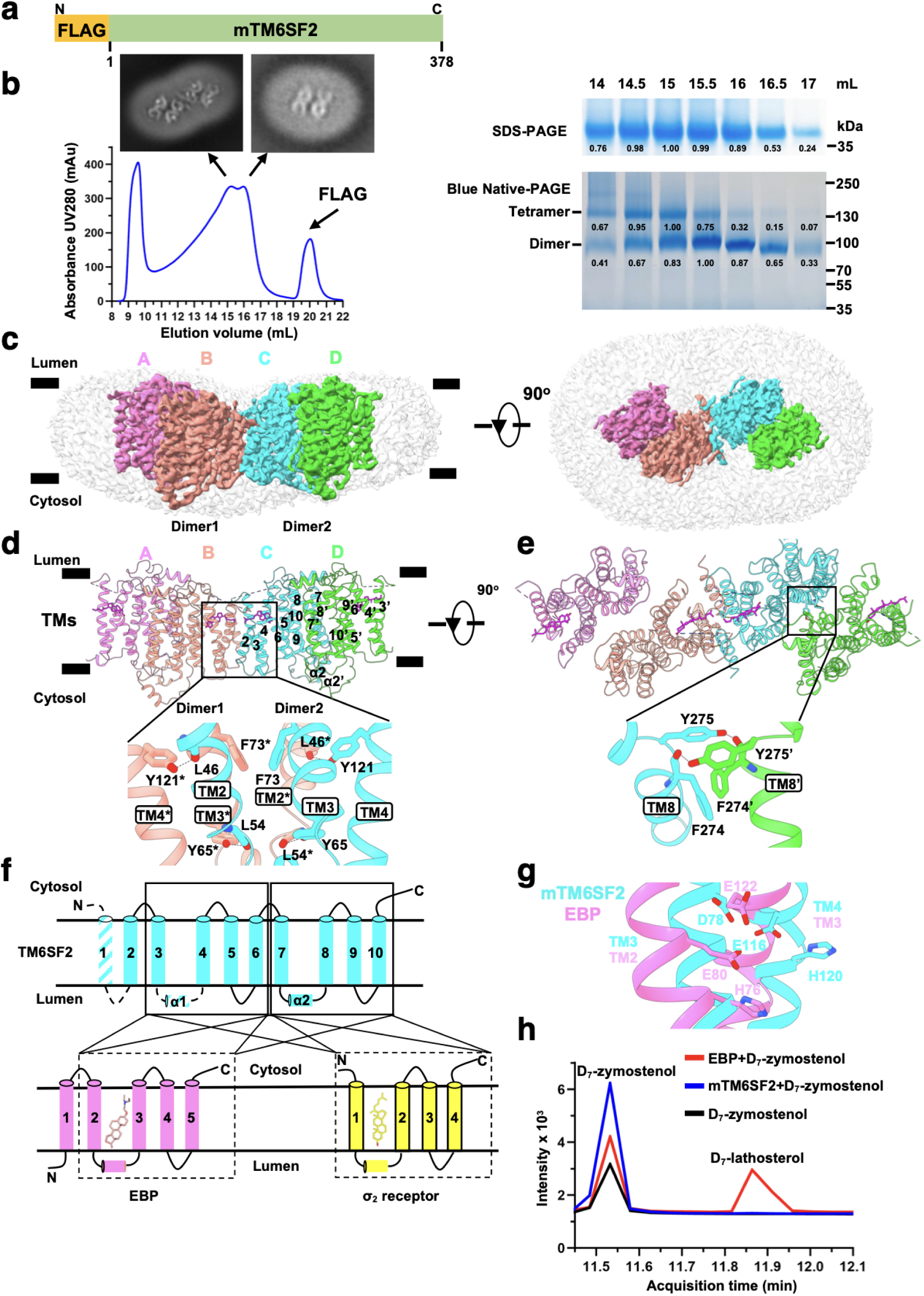
Overall structure of tetrameric TM6SF2. **a,** Schematic of the full-length mouse TM6SF2 (mTM6SF2) construct used for cryo-EM analysis. **b,** Representative size exclusion chromatogram (SEC) profile of mTM6SF2 using a Superose 6 Increase 10/300 GL column, with representative 2D class averages of tetrameric and dimeric particles shown above. Corresponding SDS-PAGE and Blue Native-PAGE gels with molecular weight markers are shown on the right. For SDS-PAGE analysis of mTM6SF2, MES running buffer was used, and mTM6SF2 migrated as a band above 35 kDa. **c,** Cryo-EM density map of the mTM6SF2 tetramer with a transparent surface showing the same map at a lower contour level (0.166) in ChimeraX. Side view of the transmembrane helical bundle (left) and bottom view (from ER lumen, right) of the four protomers colored: magenta (A), pink (B), cyan (C), and green (D). **d,** The tetrameric structure of mTM6SF2 (side view). Key interacting residues (Phe73, Tyr121, and Tyr65) at the dimer-dimer interface from protomers B and C are shown at bottom with potential hydrogen bonds indicated by dashed lines. **e,** Bottom view of the tetramer. The dimeric interface from protomers C and D is highlighted with key residues (Phe274 and Tyr275) shown at the bottom. Potential hydrogen bonds indicated by dashed lines. **f,** Schematic of shared structural features of mTM6SF2 (cyan), EBP (lower left, pink) and σ2 receptor (lower right, yellow). Disordered regions in mTM6SF2 are indicated by dashed lines. **g,** Structural alignment of mTM6SF2 (cyan) with EBP (pink). The catalytic triad of EBP (His76, Glu80, and Glu122) forming the active core is shown as sticks. Corresponding residues in mTM6SF2 (His120, Glu116, and Asp78) are highlighted, implying a putative catalytic site. **h,** mTM6SF2 does not catalyze conversion of zymostenol to lathosterol.

Initial cryo-EM analysis revealed strongly preferred orientations on holey carbon grids. After optimization of blotting and vitrification conditions, grids exhibited a gradient of ice thickness. Near the edges of the holes, where the ice was thicker, particles formed dense, strip-like assemblies, whereas at the center of the hole, where the ice was thinner, particles appeared as well-dispersed, round discs (Extended Data Fig. 1a). Data collection with a 30° stage tilt substantially improved angular sampling and yielded high-quality two-dimensional class averages (Extended Data Fig. 1b).

We determined the structure of tetrameric mTM6SF2 at 3.64 Å resolution (Fig. 1c, Extended Data Figs. 1c-e and 2a-c and Extended Data Table 1). The dimeric form of TM6SF2 could not be resolved to high resolution, likely due to its small size and increased conformational flexibility (Extended Data Fig. 3 and Extended Data Table 1). Accordingly, subsequent analyses focused on the tetramer.

The tetramer is organized as a dimer of dimers with overall C2 symmetry, which is similar to our previous findings for ACAT1 (Acetyl-CoA acetyltransferase 1), an ER enzyme that esterifies cholesterol^16^ (Fig. 1c, d). Protomers A and B form one dimer, and C and D form a second (Fig. 1c, d). Although symmetry was not imposed within dimers during cryo-EM data processing, protomers C and D adopt nearly identical conformations with a root-mean-square deviation (RMSD) of 0.530 Å (Extended Data Fig. 2d). Each protomer contains 10 TM helices with TM1 being disordered. The short α1 helix between TM3 and TM4 was not resolved, and TM2 was disordered in protomers A and D (Fig. 1d and Extended Data Figs. 2b).

The dimer-dimer interface between protomers B and C spans ∼1360 Å^2^ and is formed primarily by TM2-TM4 (Fig. 1d). Phe73 stabilizes this interface through π-π interaction, while Tyr65, Tyr121, and the main chains of Leu54 and Leu46 contribute potential hydrogen bonds (Fig. 1d). To assess the contributions of these three residues to tetramer formation, alanine was substituted for three large aromatic residues: F73, Y65 and Y121. The F73A and Y121A mutants retained tetramerization, whereas Y65A exhibited marked instability and aggregation, consistent with disruption of tetramer assembly (Extended Data Fig. 4a). Blue Native-PAGE analysis of lysates from HEK293 cells expressing mTM6SF2^Y65A^ revealed aggregates and dimers, indicating that loss of tetramerization does not preclude dimer formation (Extended Data Fig. 4b).

The dimer interface between protomers C and D spans ∼818.2 Å^2^ and involved TM8 (Fig. 1e). Phe274 stabilizes the interface via a π-π interaction, while Tyr275 and the backbone atoms of Phe274 contribute potential hydrogen bonds (Fig. 1e). Simultaneous substitution of Phe274 and Tyr275 with alanine (FY/AA) resulted in aggregation and loss of detectable dimers, consistent with destabilization of the protein (Extended Data Fig. 4a). Blue Native-PAGE analysis of HEK293 cell lysates expressing mTM6SF2^FY/AA^ confirmed extensive aggregation (Extended Data Fig. 4b).

We previously determined the structure of TM6SF1, a lysosomal protein that is a homolog of TM6SF2 and participates in mTORC1 signaling^17^. mTM6SF2 closely resembles cholesterol-bound TM6SF1 (RMSD of 0.732 Å) (Extended Data Fig. 5). Notable, TM2 in protomer C of mTM6SF2 adopts a helix-swapped arrangement, extending to interact with TM3 and TM4 of protomer B, thereby contributing to tetramer formation (Extended Data Fig.5).

Similar to TM6SF1, TM6SF2 shares structural similarity with EBP (3-β-hydroxysteroid-Δ8, Δ7-isomerase)^15,18^ and with σ2 receptor^19^ (Fig. 1f and Extended Data Fig. 6a, b). EBP is an enzyme in the cholesterol biosynthetic pathway that catalyzes conversion of zymostenol to lathosterol^15,20^, whereas σ2 receptor is an ER resident protein of unclear function^19,21^. All three proteins share EXPERA domains. TM3-TM6 and TM7-TM10 of TM6SF2 align with TM2-TM5 of EBP (RMSDs of 1.455 Å and 1.198 Å, respectively) (Fig. 1f and Extended Data Fig. 6a). Similarly, TM3-TM6 and TM7-TM10 aligns with TM1-TM4 of σ2 receptor (RMSDs of 1.154 Å and 1.155 Å, respectively) (Fig. 1f and Extended Data Fig. 6b). EBP contains a catalytic triad (His76, Glu80, and Glu122) within its conserved EXPERA domain that catalyzes conversion of zymostenol to lathosterol^15^. A corresponding triad is present in mTM6SF2 (His120, Glu116, and Asp78) (Fig. 1g), but we failed to detect any sterol isomerase activity of the protein under the conditions tested (Fig. 1h).

### TM6SF2 binds cholesterol

A density was observed within a cavity formed by TM3-TM6 in mTM6SF2, whereas no analogous pocket was evident in TM7-TM10 (Fig. 2a,b and Extended Data Fig. 6c). Structural alignment of mTM6SF2 with human TM6SF1 (hTM6SF1) revealed that the cholesterol bound to TM6SF1 localizes adjacent to the putative pocket in mTM6SF2 (Extended Data Fig. 5), suggesting that the density in mTM6SF2 corresponds to cholesterol. To test this possibility, lipids were extracted from purified mTM6SF2 and analyzed by gas chromatography-mass spectrometry (GC-MS). A peak with a retention time and fragment pattern of cholesterol was detected in samples containing mTM6SF2 (Fig. 2c). In an *in vitro* competitive binding assay, deuterated cholesterol effectively displaced unlabeled cholesterol, whereas 25-hydroxycholesterol did not (Fig. 2d), indicating specificity of binding.

**Fig. 2.**
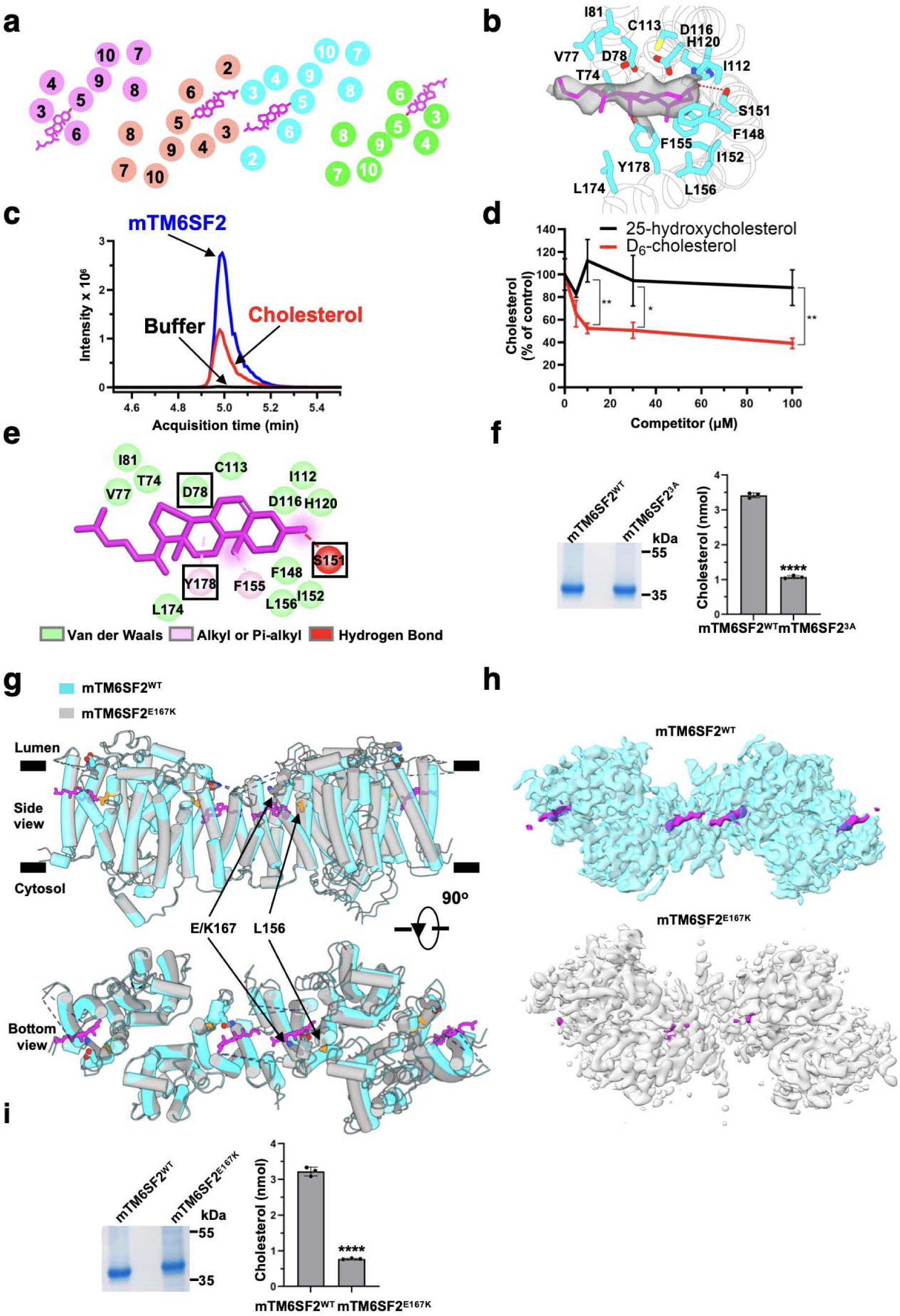
TM6SF2 binds cholesterol. **a,** Cross-sectional view of mTM6SF2 showing the TMs and the location of cholesterol molecules. **b,** Molecular interactions between mTM6SF2 and cholesterol (magenta sticks). Cryo-EM map of cholesterol is shown. Ser151 forms a potential hydrogen bond (dashed line) with the 3’-hydroxyl group of cholesterol. **c,** Liquid chromatography-mass spectrometry (LC-MS) chromatograms of sterols extracted from purified mTM6SF2 protein. Retention profiles are shown for buffer (black), Cholesterol (red), and purified mTM6SF2 (blue). The matching retention time between the cholesterol and sterol peak in the mTM6SF2 confirms direct cholesterol binding. **d,** Competitive binding of D_6_-cholesterol to mTM6SF2. Displacement of cholesterol by increasing concentrations of D_6_-cholesterol (red) or 25-hydroxycholesterol (black). Data are presented as mean ± SE (n=3); unpaired *t*-test. *P < 0.05; ** P < 0.01. **e,** Key residues involved in cholesterol binding. Glu78, Ser151, and Tyr178 are highlighted in a black box and substituted to generate the triple mutant mTM6SF2^3A^ (D78A/S151A/Y178A). **f,** Cholesterol quantification in purified mTM6SF2^WT^ and mTM6SF2^3A^. Cholesterol extracted from 2 nmol of purified mTM6SF2^WT^ or mTM6SF2^3A^ was quantified by LC-MS. Data represent mean ± SEM (n=3). **** P < 0.0001 (two-tailed *t-test*). **g,** Structural alignment of mTM6SF2 and mTM6SF2^E167K^. Side view and bottom view (from ER lumen) of superposition of mTM6SF2 (Cyan) and mTM6SF2^E167K^ (grey). Cholesterol is shown as sticks (magenta). Residues Glu167, Lys167, and Leu156 are shown as sticks. **h,** Cryo-EM density maps of mTM6SF2^WT^ (cyan) and mTM6SF2^E167K^ (grey) shown at the same contour level. Cholesterol-like densities (magenta) are visible in the wild-type but largely absent in the E167K mutant, indicating a loss of cholesterol binding. **i,** Cholesterol quantification in purified mTM6SF2^WT^ and mTM6SF2^E167K^. Cholesterol extracted from 2 nmol of proteins was measured by LC-MS. Data represent mean ± SEM (n=3). **** P < 0.0001 (Student’s *t*-test).

Structural analysis indicated that Ser151 forms a hydrogen bond with the 3’-hydroxyl group of cholesterol, while several aromatic resides, including Phe148 and Phe155 in TM5 and Tyr178 in TM6, create a hydrophobic environment that accommodates the putative sterol (Fig. 2b, e). These residues are highly conserved across species (Extended Data Fig. 7), and corresponding densities are observed at the tetramer interface (Fig. 2a, b). In addition, the side chain of Asp78 interacts with the sterol via a Van der Waal interaction (Fig. 2b, e).

To confirm cholesterol binding and assess its role in tetramer assembly, predicted sterol-coordinating residues were substituted to generate a triple mutant, (D78A/S151A/Y178A; mTM6SF2^3A^) (Fig. 2e). The amount of cholesterol in purified mTM6SF2^3A^ was ∼30% of mTM6SF2^WT^ (Fig. 2f). Blue Native-PAGE analysis showed that mTM6SF2^3A^ exists as a mixture of dimers and tetramers (Extended Data Fig. 8a, b), indicating that cholesterol binding is not required for tetramer assembly. Together, these structural and biochemical data support assignment of the cryo-EM density as cholesterol.

### E167K disrupts cholesterol binding

The E167K variant destabilizes TM6SF2 when expressed in cultured cells, consistent with the allele conferring a loss of function^1^. Glu167 is located adjacent to the cholesterol binding pocket (Fig 2g) in a well conserved region of the protein (Extended Data Fig. 7), as is another less frequent naturally-occurring missense variant (L156P) that is also associated with SLD and reduced plasma LDL-C levels^22^. To investigate the effect of the E167K substitution on TM6SF2 structure and function, we purified mTM6SF2^167K^ and analyzed the protein using SEC (Extended Data Fig. 8a, c). The yield of mTM6SF2^E167K^ was lower than that of mTM6SF2^WT^. Analysis by Blue Native-PAGE revealed a mixture of dimers and tetramers similar to mTM6SF2^WT^ (Extended Data Fig. 8a, c), indicating that the mutation did not alter the oligomeric state. Despite the reduced yield, we obtained a high-resolution structure of tetrameric mutant mTM6SF2^E167K^ using cryo-EM at 3.58 Å resolution (Fig. 2g, Extended Data Figs. 9, 10 and Extended Data Table 1). The overall conformation of mTM6SF2^E167K^ was nearly identical to that of mTM6SF2^WT^ (RMSD of 0.526 Å; Fig. 2g), indicating that the 167 substitution did not interfere with global folding of the protein.

Although the folding pattern of mTM6SF2^E167K^ was similar to the WT protein, the cholesterol-like densities were markedly reduced in the map of mTM6SF1^E167K^ compared to that of mTM6SF2^WT^ at the same contour level (Fig. 2h). LC-MS analysis showed that the amount of cholesterol bound to the mutant protein was 76% less than that seen with mTM6SF2^WT^ (Fig. 2i). We also tried to analyze the structural effects of the other naturally-occurring missense variant in TM6SF2, TM6SF2^L156P^, that is associated with increased SLD risk^22^. Unfortunately, the mutant protein aggregated during purification (Extended Data Fig. 8a).

### TM6SF2, but not TM6SF2^E167K^, binds ApoB-100

To determine if TM6SF2 interacts with ApoB-100, the major apolipoprotein of VLDL and LDL, we purified FLAG-tagged human TM6SF2 (hTM6SF2) and human LDL^23,24^ (Fig. 3a). Although we successfully purified hTM6SF2, we were unable to determine its structure by cyro-EM (Extended Data Fig. 11). Human TM6SF2 was incubated with LDL at a molar ratio of 8:1 (TM6SF2: ApoB-100). In the presence of ApoB-100, the elution volume of hTM6SF2 shifted from a peak of ∼ 16.5 mL (Fig. 3a, black solid line) to ∼11.5 mL (Fig. 3a, black dotted line), consistent with formation of an hTM6SF2-ApoB-100 complex (Fig. 3a). Quantification of SEC fractions showed that hTM6SF2 and ApoB-100 exhibited highly similar elution profiles, further supporting co-migration (Fig. 3a). These findings are consistent with prior observations^11,25^ that TM6SF2 physically interacts with ApoB.

**Fig. 3.**
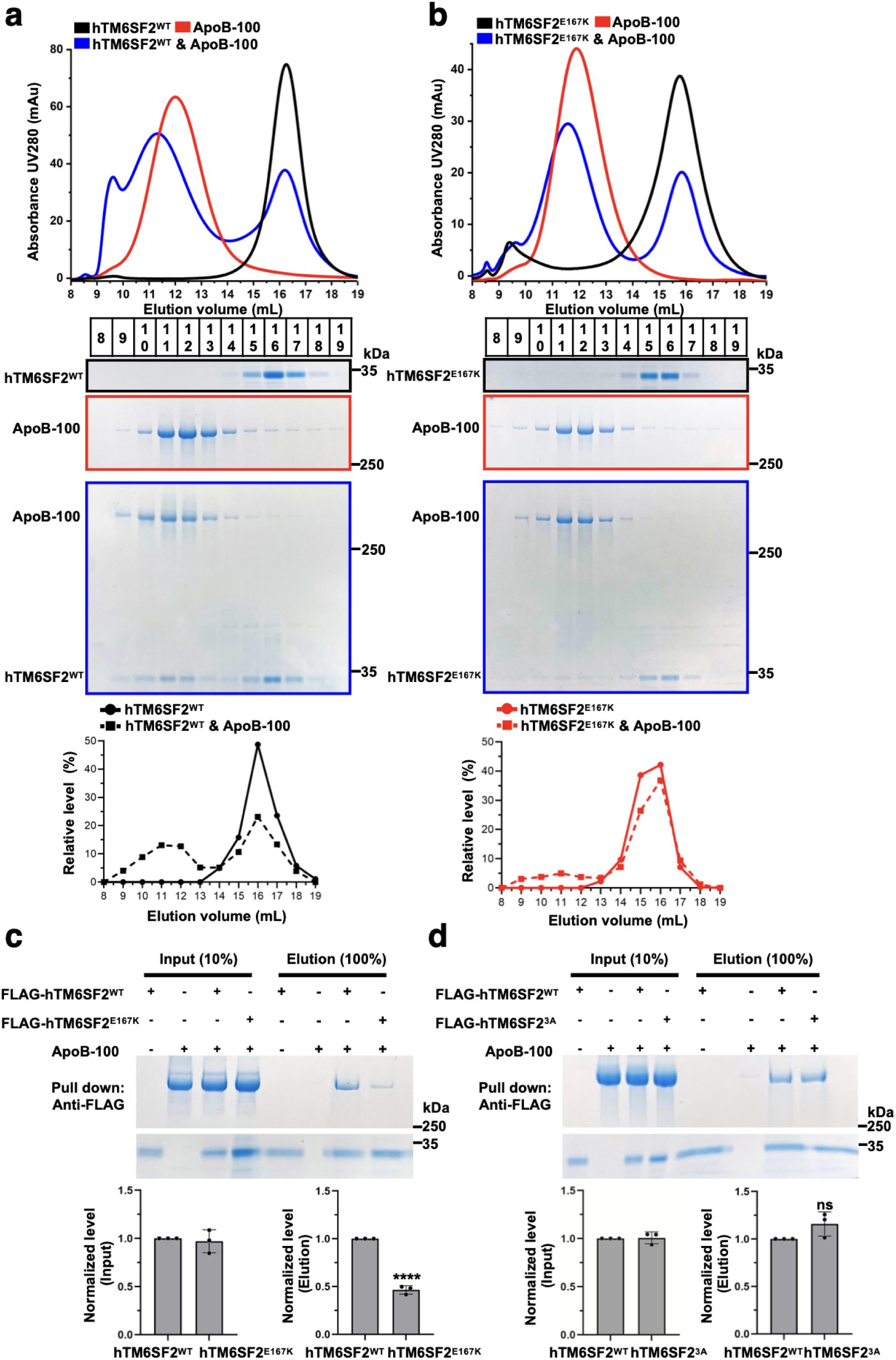
TM6SF2^E167K^ impairs ApoB-100 interaction. **a, b,** Gel filtration of hTM6SF2^WT^ **(a)** and hTM6SF2^E167K^ **(b)** in the absence or presence of ApoB-100. SDS-PAGE analysis of each fraction is shown below. The relative level of hTM6SF2^WT^ or hTM6SF2^E167K^ in each fraction before and after incubation with ApoB-100 was quantified using ImageJ and is shown at the bottom. For SDS-PAGE analysis of hTM6SF2, tris-glycine running buffer was used, and hTM6SF2 migrated as a band below 35 kDa. **c,** The E167K mutation attenuates the interaction between TM6SF2 and ApoB-100. FLAG-tagged hTM6SF2^WT^ and hTM6SF2^E167K^ were immobilized on anti-FLAG M2 resin and subjected to pulldown assays with ApoB-100. Input and elution were analyzed by 4%-12% gradient SDS-PAGE, followed by Coomassie blue staining. **d,** The cholesterol binding deficient mutant does not impair the interaction between hTM6SF2 and ApoB-100. FLAG-tagged hTM6SF2 and hTM6SF2^3A^ were immobilized on anti-FLAG M2 resin and subjected to pulldown assays with ApoB-100. In **c** and **d**, band intensities were quantified using ImageJ. Data represent mean ± SEM (n=3 independent experiments). ****p < 0.0001 (two-tailed *t-test*); ns, not significant.

To determine whether the E167K substitution alters the association of TM6SF2 with ApoB-100, we repeated the experiment using hTM6SF2^167K^. A much smaller percentage of the hTM6SF2 shifted to a higher molecular weight peak (Fig. 3b). We then performed pulldown assays to compare binding of hTM6SF2^E167K^ and hTM6SF2^WT^ to ApoB-100. Despite using comparable amounts of proteins, binding of hTM6SF2^E167K^ to ApoB-100 was reduced by ∼50% relative to hTM6SF2^WT^ (Fig. 3c). In contrast, binding of hTM6SF2^3A^ with ApoB-100 was comparable to that of hTM6SF2^WT^ (Fig. 3d), indicating that cholesterol binding is functionally independent of ApoB-100 interaction.

These data indicate that the E167K substitution alters TM6SF2 in a manner that impairs both cholesterol binding and its interaction with ApoB-containing lipoprotein. Glu167 resides in the loop connecting TM5 and TM6, adjacent to the cholesterol-binding pocket and oriented toward the ER lumen (Fig. 2g). Although substitution of Glu167 with lysine appears to disrupt cholesterol access from the lipid bilayer to the binding pocket, the reduction in cholesterol binding does not seem to account for the weakened interaction with ApoB-containing lipoproteins, since TM6SF2^3A^ bound ApoB-100 as well as TM6SF2^WT^.

### Cholesterol-bound TM6SF2 is essential for VLDL-TG secretion

We used CRISPR-Cas9 to disrupt TM6SF2 in cultured rat hepatocytes (CRL-1601 cells) (Fig. 4a, left)^26^. Disruption of TM6SF2 was confirmed by genomic DNA sequencing and by immunoblot analysis using a rabbit anti-rat TM6SF2 (rTM6SF2) polyclonal antibody that we developed^11^ (Fig. 4a, right). Nontargeted CRL-1601 cells were used as controls in all experiments.

**Fig. 4.**
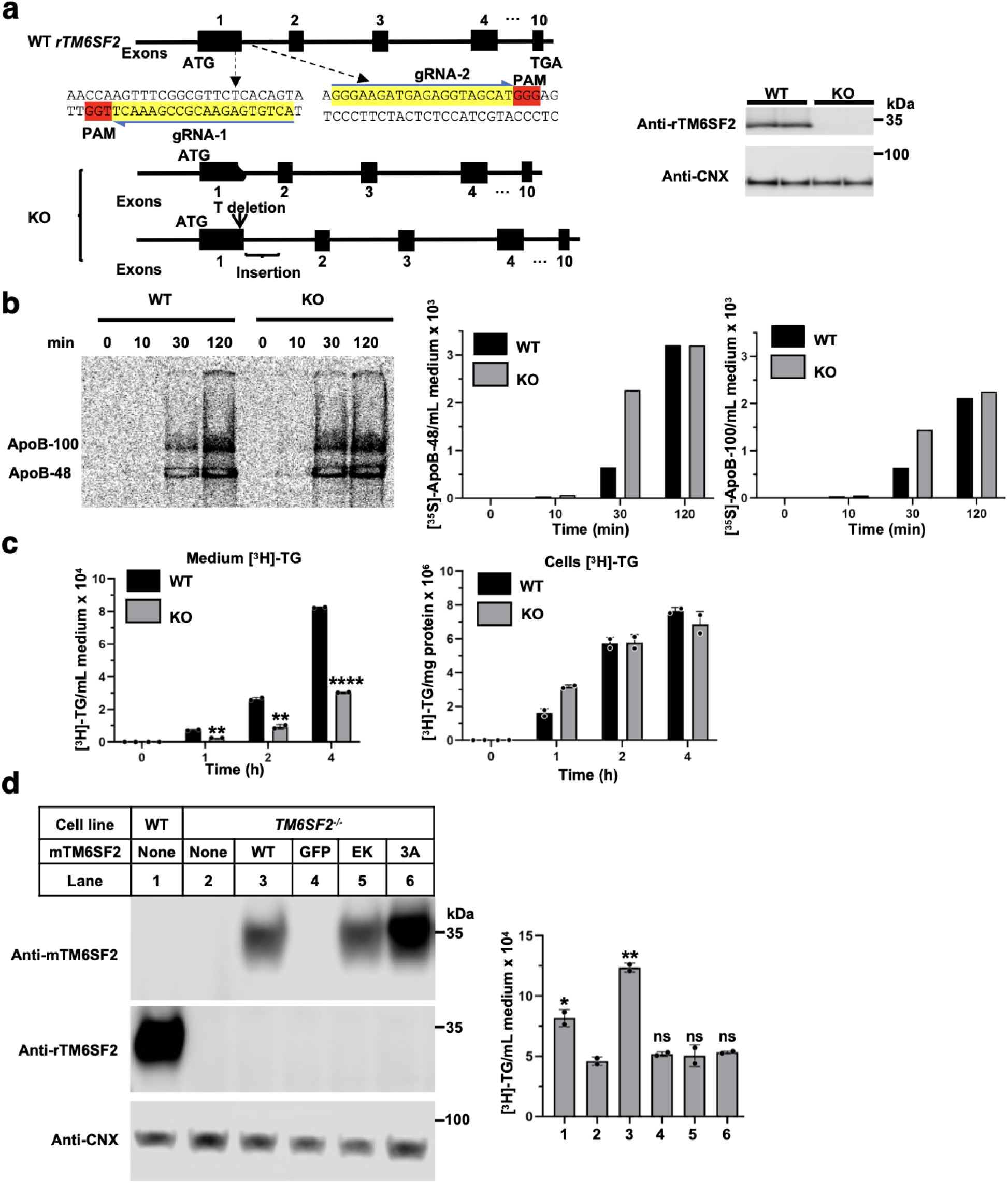
TM6SF2 is required for VLDL-TG secretion. **a,** Generation of *Tm6sf2^−/−^* CRL-1601 cells by CRISPR/Cas9. Two guide RNAs (gRNA1 and gRNA2) targeting exon 1 and the intron between exon 1 and exon 2 of rat *TM6SF2* are shown with target sequences highlighted in yellow and the PAM sequences in red. Knockouts were confirmed by immunoblotting with an anti-rat TM6SF2 antibody. Calnexin (CNX) was used as a loading control. **b,** Representative phosphor-image of radiolabeled ApoB secreted into th culture medium at the indicated chase time points. Quantification of secretion of ApoB-48 and ApoB-100 over 0-120 min chase period is shown on the right. **c,** Quantification of synthesis (left) and secretion (right) of radiolabeled triglycerides (TG) over 0-4 hours. Data are presented as mean ± SEM (n=2 independent experiments). Statistical significance was assessed by unpaired *t*-test. ** P < 0.01, **** P < 0.0001. **d,** Quantification of TG secretion. Immunoblot analysis of mouse TM6SF2 (mTM6SF2) and rat TM6SF2 (rTM6SF2) expression is shown with Calnexin (CNX) used as a loading control. Data are presented as mean ± SEM (n=2). Statistical significance was assessed by unpaired two-tailed *t*-test. * P < 0.05; ** P < 0.01; ns, not significant.

To examine the effect of TM6SF2 inactivation on ApoB-containing lipoprotein secretion, we performed pulse-chase experiments in WT and KO cells. Cells were depleted of methionine and cysteine for 1 h, followed by labeling with [^35^S]-methionine^27^ for 20 min. During the chase, media were collected at the indicated time points. The proteins were subjected to SDS-PAGE and then the gel was exposed to film. The fraction of ApoB-100 and ApoB-48 that was secreted relative to total cellular ApoB at the beginning of the experiment (cell-associated) is shown in Fig. 4b (right). KO cells showed a transient increase in ApoB-48 and ApoB-100 secretion at 30 min, which returned to WT levels by 120 min (Fig. 4b).

To assess the effect of TM6SF2 inactivation on VLDL-TG secretion, CRL-1601 cells were incubated with [^3^H]-glycerol^27^ and oleic acid, and both the cells and media were collected at the indicated time points. In contrast to ApoB-100 secretion, TG secretion was significantly reduced in KO cells at 1, 2, and 4 h; at the final timepoint, TG secretion in KO cells was reduced to ∼24-35% of WT levels (Fig. 4c, left), whereas intracellular TG levels remained unchanged (Fig. 4c, right).

For rescue experiments, we expressed mTM6SF2, which shares 92% sequence identity with rat TM6SF2 (Extended Data Fig. 7), in the *Tm6sf2^−/−^*cells (Fig.4d, left). AAV-mediated expression of mTM6SF2^WT^ restored VLDL-TG secretion in KO cells, whereas expression of mTM6SF2^E167K^ and a cholesterol-binding deficient mutant (mTM6SF2^3A^) did not (Fig. 4d, right). AAV-GFP served as a negative control in this experiment. Expression of mTM6SF2 was confirmed by immunoblotting using a mouse monoclonal antibody that we generated using a peptide from the C-terminus of mTM6SF2.

Taken together, these results support a model in which TM6SF2 promotes VLDL-TG secretion without substantially altering ApoB secretion, and cholesterol binding is required for this effect, though not for the association of ApoB with TM6SF2 (Fig. 3d).

## Discussion

In this study, we used cryo-EM to determine the structure of mouse TM6SF2. Our data showed that TM6SF2 functions as a cholesterol-bound scaffold in the smooth ER that promotes VLDL lipidation through direct interaction with ApoB-containing lipoproteins. The structure reveals a cholesterol binding pocket within the transmembrane region, and biochemical, structural, and functional analysis demonstrate that cholesterol binding is required for protein stability and for efficient VLDL-TG secretion, but not for binding to ApoB-100. These findings provide a molecular framework for understanding how TM6SF2 drives the bulk lipidation of ApoB-containing lipoproteins in the liver, the principal route for hepatic neutral lipid secretion, and how a disease-associated variant disrupts this pathway to promote SLD.

Our data inform an evolving model of VLDL synthesis and lipidation in hepatocytes^28^. During translation and translocation of ApoB into the rough ER lumen, ApoB undergoes initial lipidation, acquiring phospholipids and neutral lipids, including TG and cholesteryl esters (CE)^28,29^ by microsomal triglyceride transfer protein (MTP/PDI)^30,31^ and phospholipid transfer protein (PLTP)^32,33^, initiating formation of primordial VLDL particles^34^. These particles subsequently migrate to the smooth ER^28^, where TM6SF2 resides^11^(Fig. 5).

**Fig. 5.**
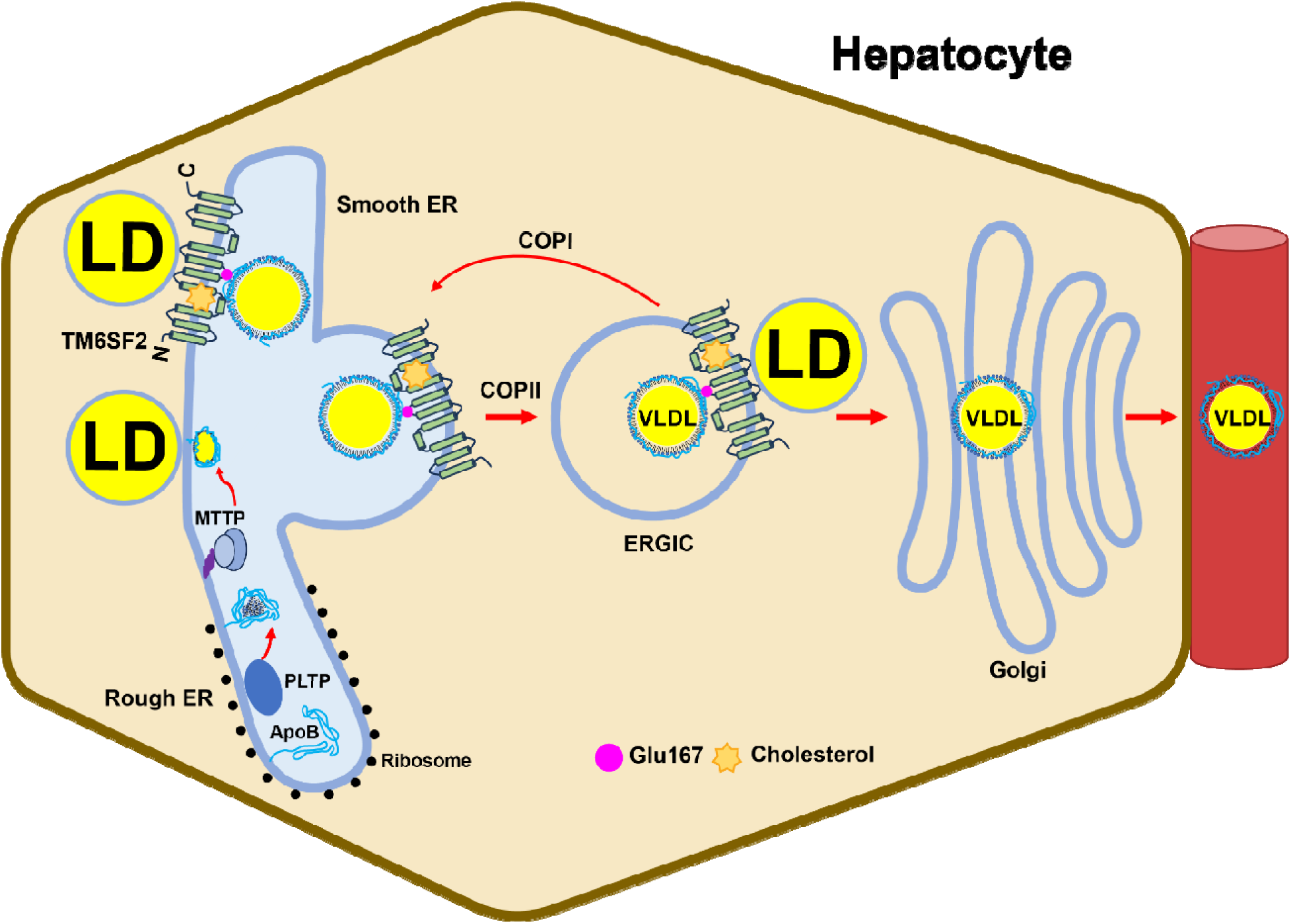
Model of VLDL lipidation mediated by cholesterol bound TM6SF2. Schematic representation of cholesterol bound TM6SF2 promoting lipoprotein interaction and recruiting lipid transfer protein (MTP/PDI, PLTP) to transfer lipids from the endoplasmic reticulum (ER) and lipid droplets (LDs), thereby forming primordial VLDL. TM6SF2 then chaperones primordial VLDL from the ER to the ER-Golgi intermediate compartment (ERGIC) and is recycled back to the ER, repeating until VLDL i fully lipidated.

TM6SF2 appears to act in the ER and ERGIC to promote bulk lipidation of VLDL, thereby converting lipid poor so-called primordial VLDL to mature, nascent TG-rich particles before secretion^34,35^. Prior studies have provided evidence that TM6SF2 and ApoB physically interact^11,25^, and suggest that TM6SF2 may function as a chaperone that facilitates trafficking of lipoproteins from the ER to the ERGIC^11^. TM6SF2 has a retrieval sequence in its cytoplasmic tail (<u>KK</u>QH)^36^ so the protein may recycle back to the ER, either alone or in association with incompletely lipidated particles^35^. We speculate that iterative cycling between the ER and ERGIC may continue until the lipoprotein acquires sufficient lipids for secretion into the circulation (Fig. 5).

Several findings from these studies support this model and establish mechanistic features of TM6SF2 function. Cholesterol binding and lipoprotein interaction represent separable and mechanistically distinct activities of TM6SF2. A cholesterol binding-deficient TM6SF2 mutant retained its interaction with ApoB-100, yet failed to restore VLDL-TG secretion (Fig. 3d, 4d), indicating that cholesterol binding is dispensable for lipoprotein association but essential for TM6SF2 function. In contrast, the E167K variant impaired both cholesterol binding and lipoprotein interaction, despite preserving the overall architecture and oligomeric state of the complex. These observations suggest that the Glu167 substitution disrupts a local environment in which cholesterol sensing is coupled to lipoprotein assembly. The molecular function of TM6SF2 and the roles of its cholesterol-binding site warrant further clarification. To date, no evidence supports TM6SF2 being directly involved in transfer of lipids to primordial VLDL, either as a transporter or transferase. Moreover, the molecular mechanism by which TM6SF2 promotes transfer of lipids to ApoB containing lipoproteins remain unclear. Our findings are most consistent with the protein providing a scaffold for the assembly and coordination of other proteins required for lipidation of VLDL, and that this function requires multimerization of the protein.

Structural analysis reveals that TM6SF2 exists as a mixture of dimers and tetramers (Fig. 1b) and disruption of oligomerization destabilized TM6SF2 (Extended Data Fig. 4a, b), supporting a functional role for higher order assembly in lipoprotein biogenesis.

Although TM6SF2 and TM6SF1 share sequence homology and structural similarity, including a conserved cholesterol binding pocket (Extended Data Fig. 5a, b), their functions diverge. TM6SF2 resides in the ER, where it interacts with ApoB-containing lipoproteins to promote the lipidation of VLDL, whereas TM6SF1 localizes to lysosomes and interacts with LAMTOR1 in the Ragulator complex to regulate mTORC1 signaling^17^. Structurally, TM1 in TM6SF2 is disordered and TM2 from one protomer undergoes a helix-swapping arrangement to interact with TM3 and TM4 of a neighboring protomer, thereby promoting tetramer formation. In contrast, TM1 in TM6SF1 sterically occludes such interactions, restricting the protein to a dimeric state (Extended Data Fig. 5a, b).

Our findings provided new mechanistic insight into TM6SF2-associated human disease; TM6SF2^E167K^, and another TM6SF2 variant, TM6SF2^L156P^ ^22^ are both associated with hepatic TG accumulation and reduced plasma LDL-C levels. Both variants are positioned adjacent to the cholesterol binding pocket. The E167K substitution interferes with both cholesterol binding and ApoB interaction (Fig. 2i and 3 b, c). The L156P is also predicted to interfere with cholesterol binding through hydrophobic interactions (Fig 2b, e), but it appears to also interfere in a more global manner with the folding of the protein (Extended Data Fig. 8a).

PNPLA3^I148M^ ^10^and TM6SF2^E167K 1^ both contribute to SLD by distinct mechanisms yet both variants confer a similar risk of hepatic steatosis and progression to advanced liver disease.^4,37^ PNPLA3^I148M^ is a gain-of-function mutation that promotes hepatic steatosis by inhibiting ATGL-mediated triglyceride hydrolysis in mouse models^38^, whereas TM6SF2^E167K^ is a loss-of-function mutation that destabilizes the protein and impairs both cholesterol binding and lipoprotein interaction, resulting in increased hepatic TG accumulation. However, in contrast to PNPLA3^I148M^, TM6SF2^E167K^ is also associated with reduced plasma LDL-C and a lower risk of CAD^2^, thus uncoupling SLD from CAD.

Our results have implications for the development of therapies to prevent SLD in carriers with TM6SF2^E167K^, and possibly for other disease-causing mutations. Previously, we found that this variant markedly reduced expression of the protein^10^. We propose that E167K has multiple effects on TM6SF2 that contribute to loss of its function: reduced protein abundance, impaired cholesterol binding, and a diminished interaction with ApoB-100. We were therefore surprised that the structure of TM6SF2^E167K^ was superimposable with that of the WT protein (Fig. 2g). This observation suggests that searching for chemical chaperones to enhance folding of the mutant protein, as has been done for several missense variant in other proteins^39^, is unlikely to be successful.

## Method

### Protein expression and purification of TM6SF2

Complementary DNAs encoding human TM6SF2 (hTM6SF2, NCBI reference sequence: NM_001001524.3) and mouse TM6SF2 (mTM6SF2, NM_001293795.1) were cloned into the pEG BacMam^40^ with an N-terminal FLAG tag. Site-directed mutations were introduced into the coding regions of hTM6SF2 or mTM6SF2 using a two-step PCR method. The coding sequences were validated by Sanger sequencing. Proteins were expressed in HEK 293 GnTI^−^ cells (ATCC, CRL-3022) using the baculovirus system. Baculoviruses were generated by transfecting *Sf9* cells with Bacmid DNA using Cellfectin™ II Reagent (Thermo Fisher Scientific, 10362100). After two rounds of amplification, the resulting viral stocks were used to infect HEK 293 GnTI^−^ cells at a density of 4×10^6^ cells/mL. After 8 hours at 37, sodium butyrate was added to a final concentration of 10 mM and the temperature was reduced to 30. After 72 hours, cells were homogenized in buffer A (20 mM HEPES, PH 7.5, 150 mM NaCl) supplemented with proteases inhibitor cocktail (Roche), leupeptin (10 μg/mL) and PMSF (1 mM) and then lysed by sonication. Cell debris was removed by low-speed centrifugation, and the supernatant was incubated with 1% (w/v) Glyco-diosgenin (GDN) for 1 hour at 4. The insoluble fraction was removed by centrifugation (15,000 rpm, 4, 30 min) and the supernatant was loaded onto an anti-FLAG M2 resin (Sigma) that had been pre-equilibrated with buffer B (buffer A plus 0.02% GDN). The resin was washed sequentially with 15 ml of Buffer B by gravity flow. The target protein was eluted in 7.5 mL of buffer C (buffer B supplemented with 0.1 mg/mL 3×FLAG peptide) and purified by size-exclusion chromatography (SEC) using a Superose 6 Increase 10/300 GL column pre-equilibrated with buffer B. The purified samples were stored at −80 °C for future analysis. For cryo-EM sample preparation, peak fractions containing the mTM6SF2 were concentrated and subjected to a second round of SEC using the same column, pre-equilibrated with buffer B. The final peak fractions corresponding to the tetramer or dimer of mTM6SF2 were concentrated and used for cryo-EM grid preparation.

### Cryo-EM sample preparation and data collection

A total of 3 μL of purified protein (10 mg/mL) was applied to glow-discharged Quantifoil R1.2/1.3 400 mesh Au holey carbon grids (Quantifoil). Grids were blotted using a Vitrobot Mark (FEI) with a blotting force of −5, blotting time of 4.5 s, at 100% humidity at 22 and then plunged into liquid ethane cooled by liquid nitrogen. For mTM6SF2 (tetramer), mTM6SF2 (dimer), and mTM6SF2^E167K^ (tetramer), a total of 9873, 5040, and 9598 raw movie stacks, respectively, were collected using SerialEM on a 300 kV Titan Krios (FEI) equipped with a K3 detector. Data were acquired at a physical pixel size of 0.827 Å per pixel and a nominal magnification of 105,000 ×, with the stage tilted at 30°. The defocus range was set to 0.8∼1.8 μm. Each movie stack was recorded over 5 s and dose-fractionated into 50 frames with a total dose of ∼60 electrons per A^2^ and an energy filter slit width 20 eV.

### Cryo-EM data processing

Raw movie stacks (0.827 Å per pixel) were gain-normalized. Motion correction for beam-induced motion was performed using MotionCor2^41^ and the contrast transfer function (CTF) was estimated using CTFFIND4^42^ within Relion 3.1.4^43^. For mTM6SF2 (tetramer), mTM6SF2 (dimer), and mTM6SF2^E167K^ (tetramer), A total of 1,635,528, 2,018,819, and 1,420,708 particles, respectively, were automatically picked using crYOLO-v1.9.7^44^ with the general model and a particle threshold of 0.3. Particles extractions were carried out in RELION3.1.4 using a box size of 350 pixels. Subsequent 2D classification, *ab initio* reconstruction, heterogeneous refinement, non-uniform refinement and local refinement was performed in CryoSPARC v4^45^.

### Model building and refinement

The initial model was built *de novo* by AlphaFold2 and docked into the cryo-EM map using ChimeraX^46^. Manual model adjustments were performed in COOT^47^, followed by real-space refinement in Phenix^48^. Structural model validation was carried out using Phenix and MolProbity^49^, with the results summarized in Extended Data Table 1. All structural figures were prepared using ChimeraX and PyMOL (www.pymol.org).

### ApoB-100 purification

Human LDL (Thermo Fisher Scientific, L3486) was filtered through a 0.22 μm membrane and further purified by SEC using a Superose 6 Increase 10/300 GL column pre-equilibrated with buffer A. Peak fractions containing ApoB-100 were pooled, concentrated, and subjected to a second round of SEC using buffer B. The peak fractions were analyzed by 4%-12% gradient SDS-PAGE (Thermo Fisher Scientific, XP04125BOX) using Tris-glycine buffer, followed by Coomassie blue staining (Abcam, ab119211). Fractions containing ApoB-100 on LDL were pooled and concentrated for subsequent analysis.

### Pulldown assay

Purified hTM6SF2 or mutant proteins and ApoB-100 were mixed at a 4:1 molar ratio and incubated on ice for 1 h. Subsequently, 20 µL of a 50% suspension of anti-FLAG M2 magnetic beads (Sigma, M8823) was added to the mixture and incubated for an additional 1 hour at 4°C. Beads were then washed three times with buffer B and eluted with buffer C. Eluted proteins were separated by SDS-PAGE (Thermo Fisher Scientific, XP04125BOX) and visualized by Coomassie blue staining (Abcam, ab119211).

### Gel-filtration analysis

Purified hTM6SF2 and ApoB-100 were mixed at an 8:1 molar ratio and then incubated on ice for 1 h. The mixture after the incubation was subjected to size-exclusion chromatography (SEC) using a Superose 6 Increase 10/300 GL column pre-equilibrated with buffer B. Eluted fractions were analyzed by SDS-PAGE (Thermo Fisher Scientific, XP04125BOX) using Tris-glycine buffer and visualized by Coomassie blue staining (Abcam, ab119211).

### Generation of TM6SF2 knockout cells

TM6SF2 knockout (KO) CRL-1601 rat hepatocytes were generated using CRISPR-Cas9 technology^26^. Two guide RNAs targeting exon 1 and the intron between exon1 and exon 2 of the rat TM6SF2 gene were designed (gRNA1: 5’-ACTGTGAGAACGCCGAAACT-3’; gRNA2: 5’-GGGAAGATGAGAGGTAGCAT-3’). Annealed oligonucleotides were cloned into pX459Neo plasmid and transfected into CRL-1601 cells using Lipofectamine™ 3000 Transfection Reagent (Invitrogen, L3000008), followed by Neomycin selection. Single-cell clones were isolated by limiting dilution, expanded, and screened for TM6SF2 expression by immunoblotting using a polyclonal rabbit anti-rat TM6SF2 antibody (505E)^11^, with Calnexin (CNX) (Enzo, ADI-SPA-860-F) as a loading control. Clones lacking TM6SF2 expression after at least three passages were designated TM6SF2 KO cell lines.

### Generation of a monoclonal antibody against mouse TM6SF2

The mouse monoclonal anti-mouse TM6SF2 antibody (9E9, lgG) was generated by fusing SP2-mIL6 mouse myeloma cells (ATCC, CRL-2016) with splenic B lymphocytes from immunized mice, as previously described^50^. Animal procedures were approved by the UT southwestern Institutional Animal Care and Research Advisory Committee (protocol no.2017-102391). Briefly, 6-8 weeks old male New Zealand Black (NZBWF1/J) mice (Jackson Laboratory, 100008) were immunized with purified recombinant C-terminal residues 291-376 of mouse TM6SF2 (50 μg), expressed in *E. coli* under denaturing conditions (8 M urea), and emulsified with Sigma Adjuvant system. Mice received one primary immunization followed by twelve booster immunizations. Hybridoma supernatants were screened by ELISA and immunoblotting, and clone 9E9 was selected for its specific recognition of mTM6SF2.

### Production of adeno-associated virus (AAV)

cDNAs encoding mTM6SF2 or its mutants or GFP were cloned into the pAAVsc-pTK vector, which was generated by inserting the pTK cassette into the pAAVsc-TBG vectors (University of Massachusetts). All coding sequences were validated by Sanger sequencing. Recombinant AAV particles were produced by co-transfecting HEK 293 cells with three plasmids: pAAVsc-pTK constructs encoding mTM6SF2 or its mutants (or GFP), pAAV-DJ (Cell Biolabs 340001), and pHelper (Cell Biolabs, 340202). Viral particles were subsequently purified according to the manufacturer’s instructions. Viral titers were determined by quantitative PCR (qPCR) using primers (forward: 5’-GGAACCCCTAGTGATGGAGTT-3’; reverse: 5’-CGGCCTCAGTGAGCGA-3’) to estimate viral genome copies. The resulting AAV were used for expression studies.

### VLDL-TG and ApoB secretion assay

VLDL-TG secretion was measured as described previously^27^. For basal secretion, on day 0, WT and TM6SF2 KO CRL-1601 cells were set up in Dulbecco’s modified Eagle’s medium (DMEM)-high glucose supplemented with 10% fetal calf serum (FCS) and penicillin-streptomycin (P/S) at a density of 2.5 × 10^5^ cells per well. On day 2, cells were washed once with PBS and incubated with 1 ml of labeling medium containing [^3^H]-glycerol (5 μCi mL^−1^) and 0.4 mM oleic acid (OA) in DMEM supplemented with 10% FCS for 0, 1, 2, and 4 hours, at each time point, culture medium were collected, and cells were washed with three time of PBS.

For rescue assay, on day 0, WT and TM6SF2 KO CRL-1601 cells were seeded under the same conditions. On day 1, cells were infected with AAV vectors expressing wild-type mTM6SF2, mTM6SF2 mutants, or GFP as a negative control. On day 3, Cells were split and reseeded at a density of 4.0 × 10^5^ cells/well. On day 5, cells were washed and incubated with [^3^H]-glycerol (5 μCi mL^−1^) and 0.4 mM OA in DMEM supplemented with 10% FCS for 3 hours. Cellular lipids were extracted using 5 mL of chloroform: methanol (2:1, v/v) and 1 mL PBS at room temperature. For culture media, 5 mL of chloroform: methanol (2:1, v/v) and 1 mL of PBS were added, followed by centrifugation at 3,000 rpm for 5 min to induce phase separation. The lower organic phase was collected and dried under nitrogen gas. Dried lipid extracts were resuspended in chloroform: methanol (2:1, v/v) and separated by thin-layer chromatography (TLC) using a solvent system of hexane: ethyl ether: acetic acid (120:30:3, v/v). TLC plate was dried and stained with iodine vapor, and triglyceride (TG) bands were scraped and quantified by scintillation counting using a liquid scintillation analyzer (Tri-Carb 2800TR).

For ApoB secretion assay^27^, cells were prepared as above. On day 2, cells were pre-incubated in methionine and cysteine free (MCF) medium for 1 hour prior to labeling. For pulse labeling, cells were incubated with [^35^S]-methionine (150 μCi mL^−1^) for 20 min at 37°C. Following labeling, cells were washed and incubated in DMEM supplemented with 10% FCS for chase periods of 0, 10, 30, and 120 min. At each time point, culture media were collected, and ApoB was immunoprecipitated using anti-ApoB antibody (Abcam, ab20737). Immunoprecipitated samples were resolved by SDS-PAGE. Gels were fixed, treated with an amplification reagent, and exposed for 4 h. Radiolabeled ApoB was detected using a Typhoon phosphorimager (Cytiva) and quantified accordingly.

### Enzymatic activity analysis in vitro

Enzymatic activity was measured as described previously^17^. Briefly, 2 nmol of purified mTM6SF2 protein in buffer B was incubated with 50 μM D_7_-zymostenol (Avanti, 700117) at 37°C for 12 hours in a final reaction volume of 200 μL. Purified EBP protein^15^ was used as a positive control, and buffer B alone served as a negative control. Reactions were quenched by addition of 10 μL chloroform/methanol (2:1, v/v). Sterol extraction, derivatization, and gas chromatography-mass spectrometry (GC-MS) analysis were performed as described previously^51^. Substrate and product were analyzed in selected ion-monitoring (SIM) mode.

### Identification and quantification of sterols

Sterols were extracted and analyzed by liquid chromatography-mass spectrometry (LC-MS) as described previously^52^. Briefly, 2 nmol of purified protein, cholesterol, or buffer control was transferred into 16 mm glass tubes and brought to a final volume of 1 mL with PBS. Methanol (1 mL) and methyl tert-butyl ether (MTBE, 2 mL) were added, followed by vortexing and centrifugation at 3,500 rpm for 5 min to induce phase separation. Approximately two-thirds of the upper organic phase was transferred to a new tube. The extraction was repeated by adding 2 mL MTBE to the original tube, and the organic phases were combined. Samples were dried under a stream of nitrogen at ∼40.

For saponification, 1 mL of 0.5 M KOH in methanol was added, and samples were incubated at 80 for 1 hour, followed by cooling to room temperature. Methanol (1 mL) and MTBE (2 mL) were added, and the extraction procedure was repeated as described above. The combined organic phases were dried under nitrogen at ∼40. Dried extracts were resuspended in 300 μL of 90% methanol and incubated at room temperature for 1 hour. Samples were transferred to 2 mL autosampler vials containing 0.5 mL flatbottom silanized inserts preloaded with 20 μL of D_7_-cholesterol (Avanti, LM4100) as an internal standard and analyzed by LC-MS. Sterols were quantified using internal standard normalization.

### D_6_-Cholesterol competitive binding assays

Each reaction contained 2 nmol of purified mTM6SF2 protein, supplemented with varying concentrations of competitor compounds [25-hydroxycholesterol (Sigma, H1015) or D_6_-cholesterol (Avanti, 700172)], and brought to a final volume of 200 μL with buffer B. The mixtures were incubated for 1 hour at 4°C. Subsequently, 40 μL of a 50% suspension of Anti-FLAG M2 Magnetic Beads (Sigma, M8823), pre-equilibrated with buffer B, was added to each reaction and incubated for an additional hour at 4°C. The beads were then washed three times with buffer B using a DynaMag-2 magnetic rack (Invitrogen, 12321D). FLAG-tagged proteins were eluted with 200 μL buffer C and the eluates were analyzed for sterols by LC-MS using 2,3,4-¹³C - cholesterol (Cambridge Isotope Laboratories, CLM-9139-0.005) as an internal standard. Cholesterol was quantified using internal standard normalization.

## Data, code, and materials availability

All data are available in the main text or the supplementary materials. The cryo-EM density maps of tetrameric mTM6SF2, dimeric mTM6SF2 and tetrameric mTM6SF2^E167K^ have been deposited in the Electron Microscopy Data Bank (EMDB) under accession codes EMD-78328, EMD-78329 and EMD-78330, respectively. The corresponding atomic coordinates of tetrameric mTM6SF2 and tetrameric mTM6SF2^E167K^ have been deposited in the Protein Data Bank (PDB) under accession codes 37NP and 37NQ, respectively.

## Acknowledgements

The cryo-electron microscopy (cryo-EM) data were collected at the University of Texas Southwestern Medical Center (UTSW) cryo-EM Facility and Howard Hughes Medical Institute Janelia cryo-EM Facility. We would like to thank Lisa Kinch and Xiao-Song Xie for helpful discussions, Eriks Smagris and Fang Xu for excellent technical support. We thank Lisa Beatty for cell culture work and Linda Donnelly for antibody generation. We thank Jeffrey McDonald of the Lipid Mass Spectrometry Core and Andrew Lemoff in the Proteomics Core, both at UTSW, for measurement of tissue lipid and protein quantification, respectively. This work was supported by NIH-R01DK090066, P30DK127984, P01-HL160487, R35 GM149533, Welch Foundation (I-1957) and American Heart Association (23EIA1038669).

## Author Contributions

S.H., X.L. and H.H.H., designed the experiments; S.H. determined the structures; S.H., J.W., and M.A.M. conducted the functional experiments; all the authors analyzed data and S.H., J.C.C., X.L. and H.H.H. wrote the paper.

---

**Extended Data Fig. 1.**
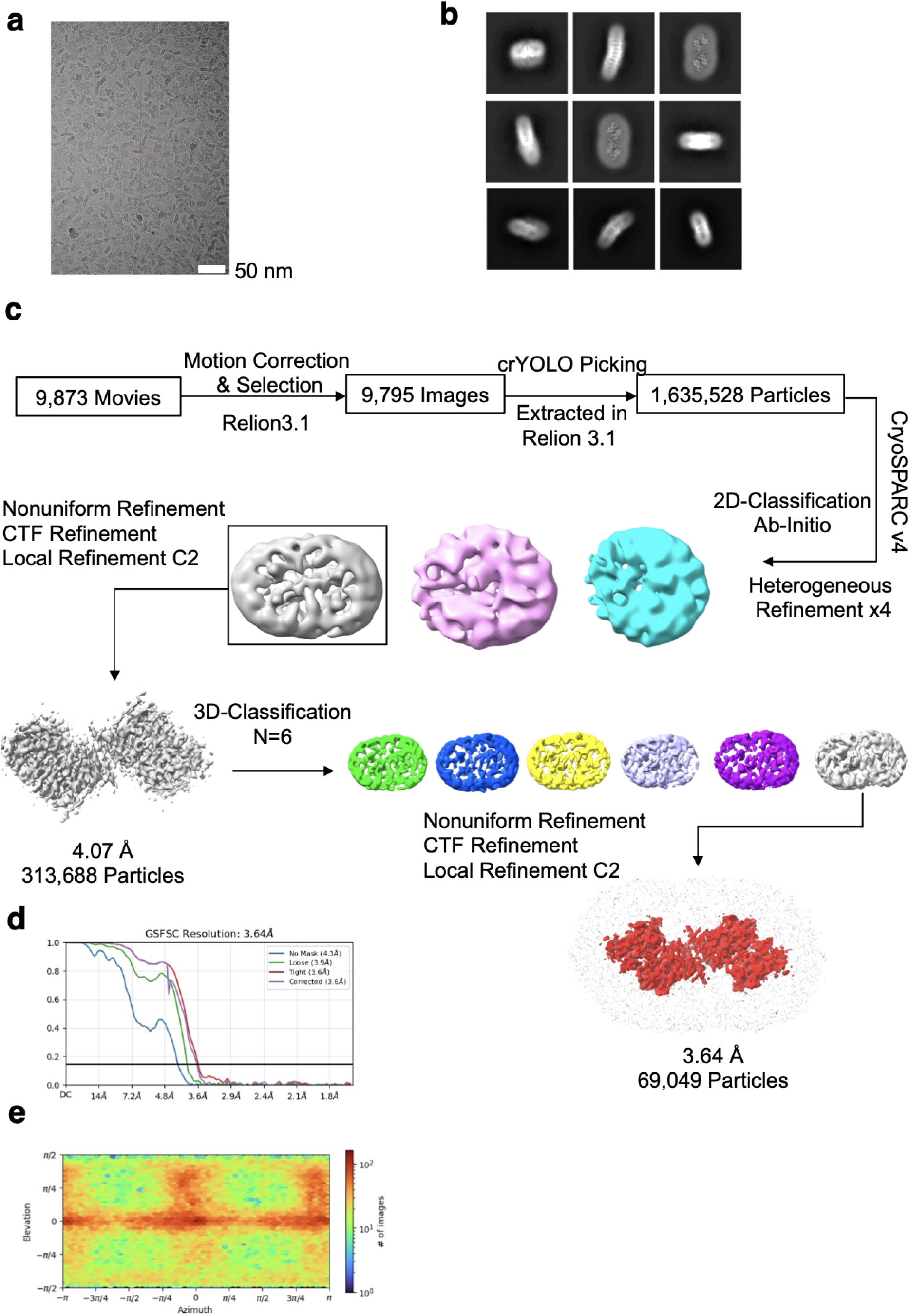
Cryo-EM analysis of tetrameric mTM6SF2. **a,** Representative cryo-EM micrograph of tetrameric mTM6SF2 in GDN micelles. Scale bar, 50 nm. **b,** Representative 2D class averages of tetramer with box size of 289.45 Å. **c,** Workflow of image processing for the mTM6SF2 tetramer. **d,** Fourier shell correlation (FSC) curves between two half maps of the mTM6SF2 tetramer. **e,** Angular distribution of particles used in the final 3D reconstruction of mTM6SF2 tetramer.

**Extended Data Fig. 2.**
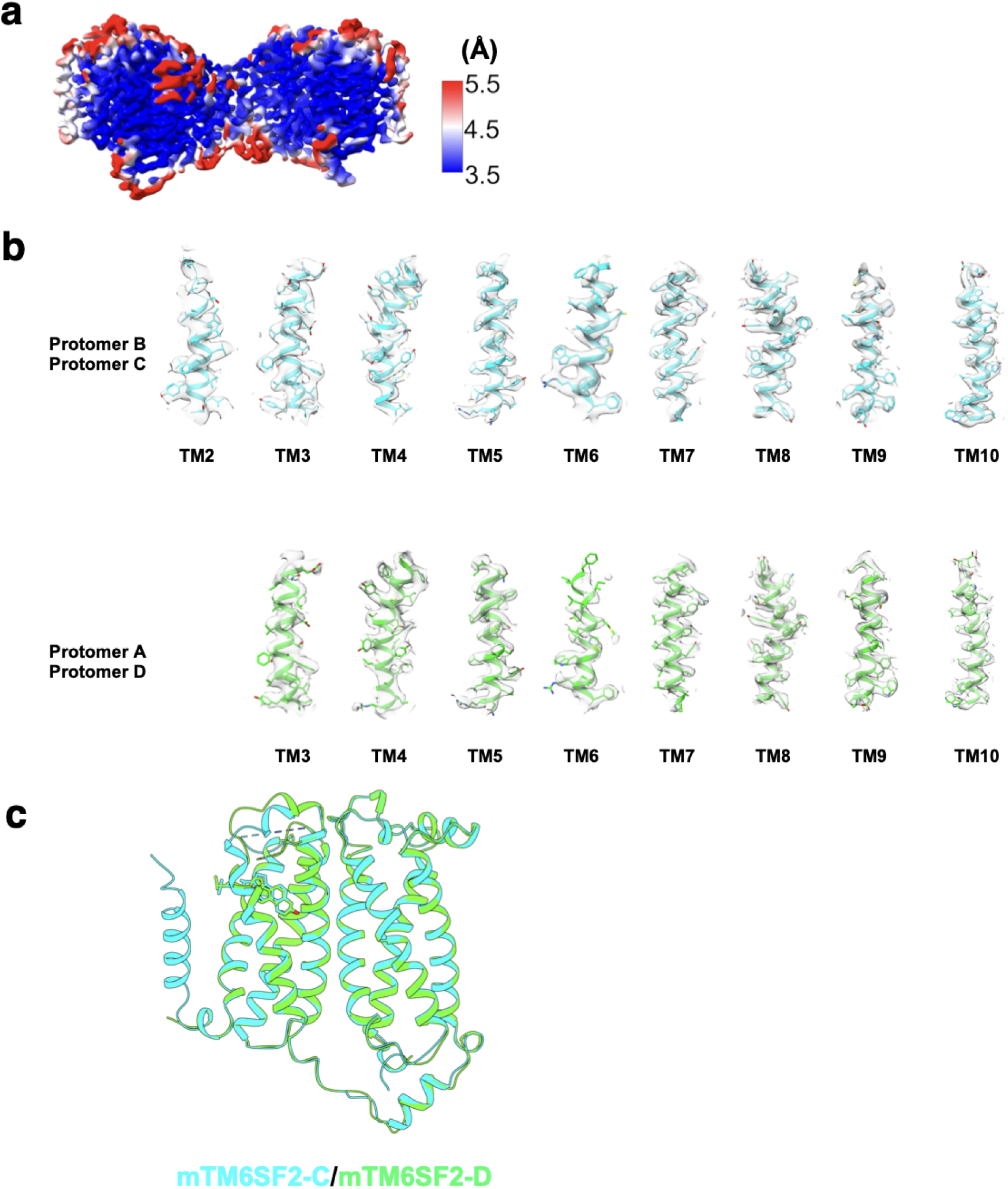
Cryo-EM density maps of tetrameric mTM6SF2. **a,** Local resolution map of the mTM6SF2 tetramer, colored according to local resolution estimated by cryoSPARC. The color gradient (blue-white-red) represents a resolution range from 3.5 Å to 5.5 Å. Cryo-EM density of the transmembrane (TM) helices of **b,** protomer B and C and **c,** protomer A and D. **d,** Structural superposition of two protomers within the dimer, showing nearly identical conformations.

**Extended Data Fig. 3.**
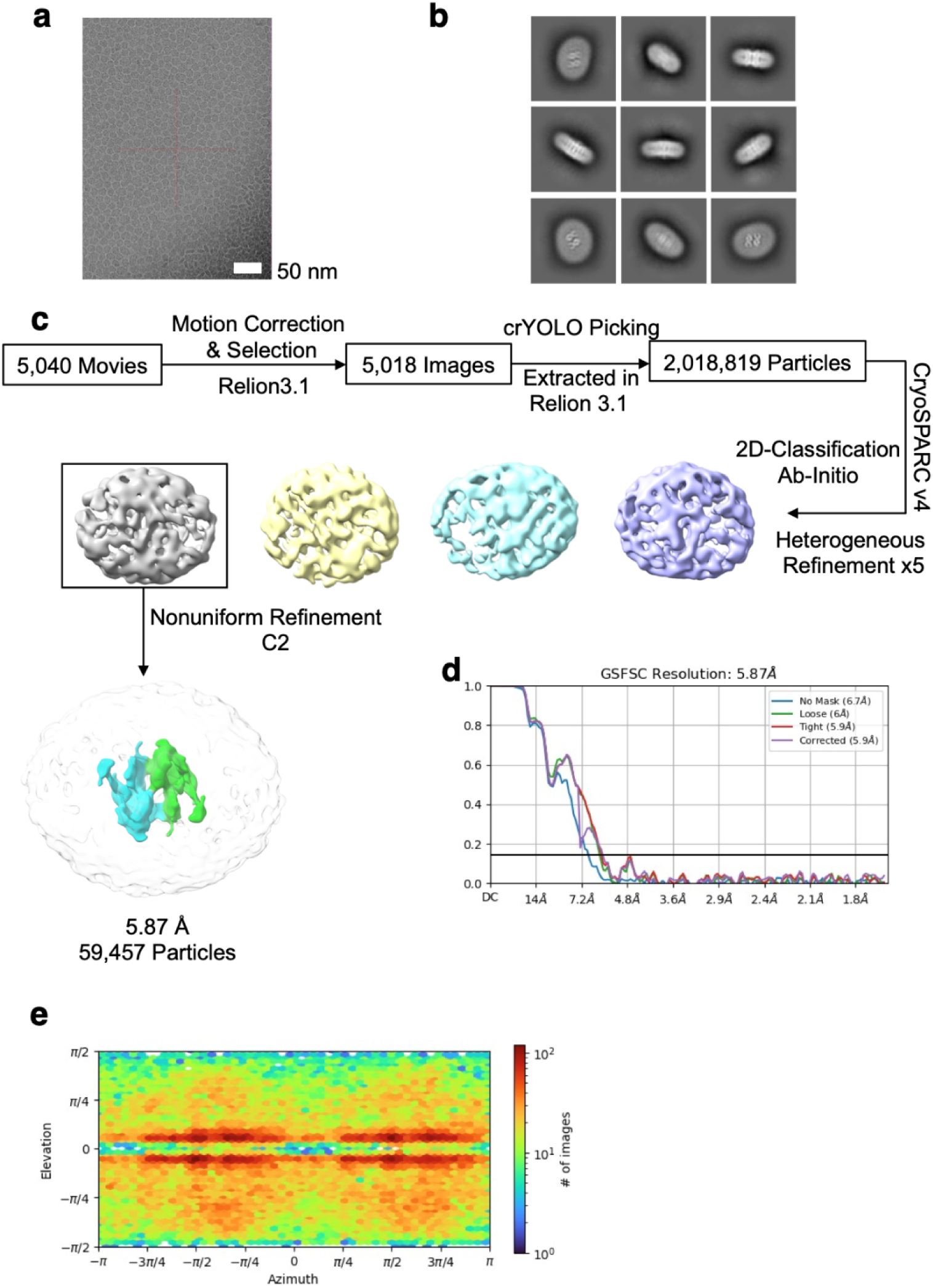
Cryo-EM analyses of dimeric mTM6SF2. **a,** Representative cryo-EM micrograph of dimeric mTM6SF2 in GDN micelles. Scale bar, 50 nm. **b,** Representative 2D class averages of dimer (box size, 289.45 Å). **c,** Workflow of image processing for the mTM6SF2 dimer. **d,** Fourier shell correlation (FSC) curves between two half maps of the mTM6SF2 dimer. **e,** Angular distribution of particles used in the final 3D reconstruction of mTM6SF2 dimer.

**Extended Data Fig. 4.**
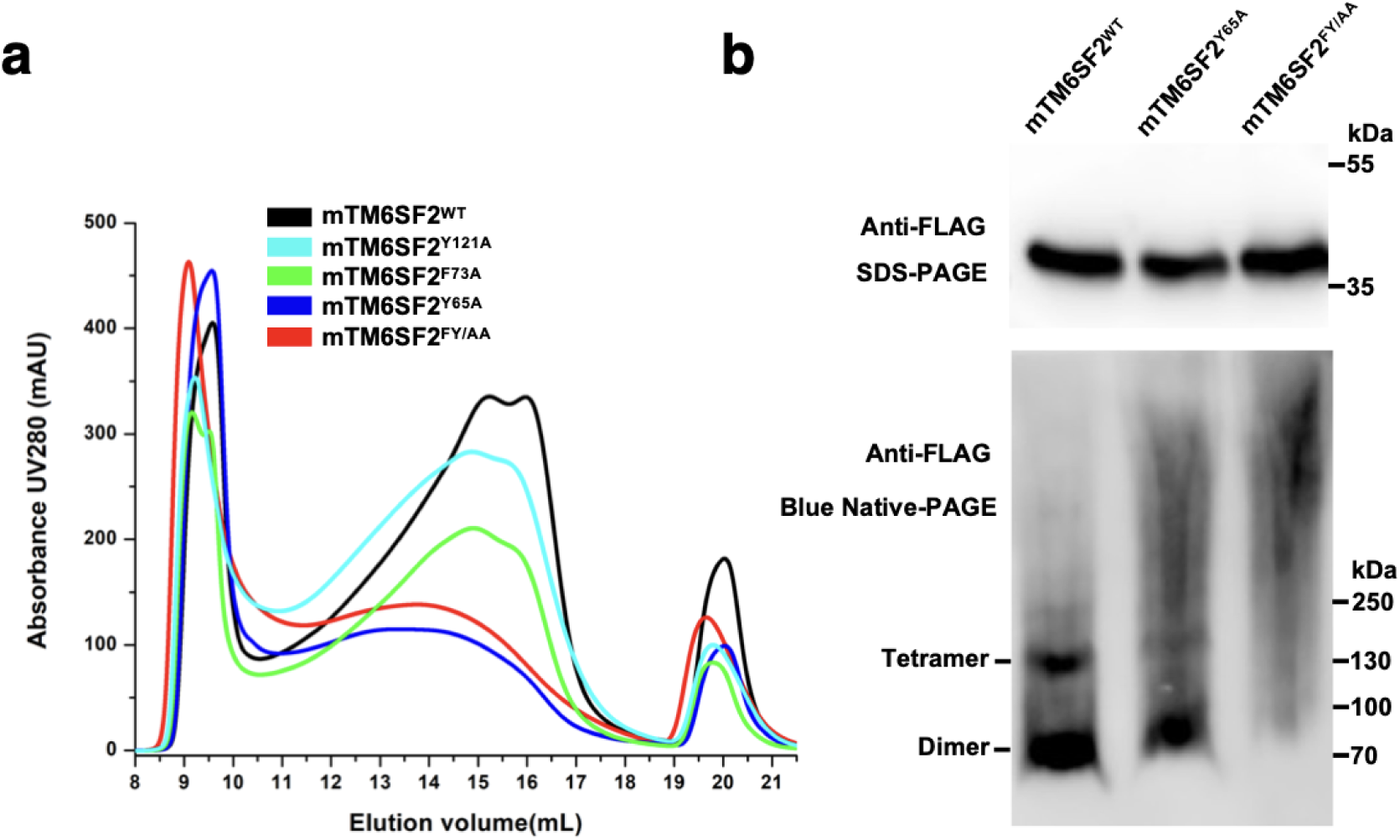
Oligomeric interface analysis. **a,** Representative SEC profiles of mTM6SF2 oligomeric interface mutants. **b,** SDS-PAGE and Blue Native-PAGE analysis of HEK293 cell lysates expressing FLAG-tagged mTM6SF2^WT^, mTM6SF2^Y65A^, and mTM6SF2^FY/AA^. Expression was detected using an anti-FLAG antibody.

**Extended Data Fig. 5.**
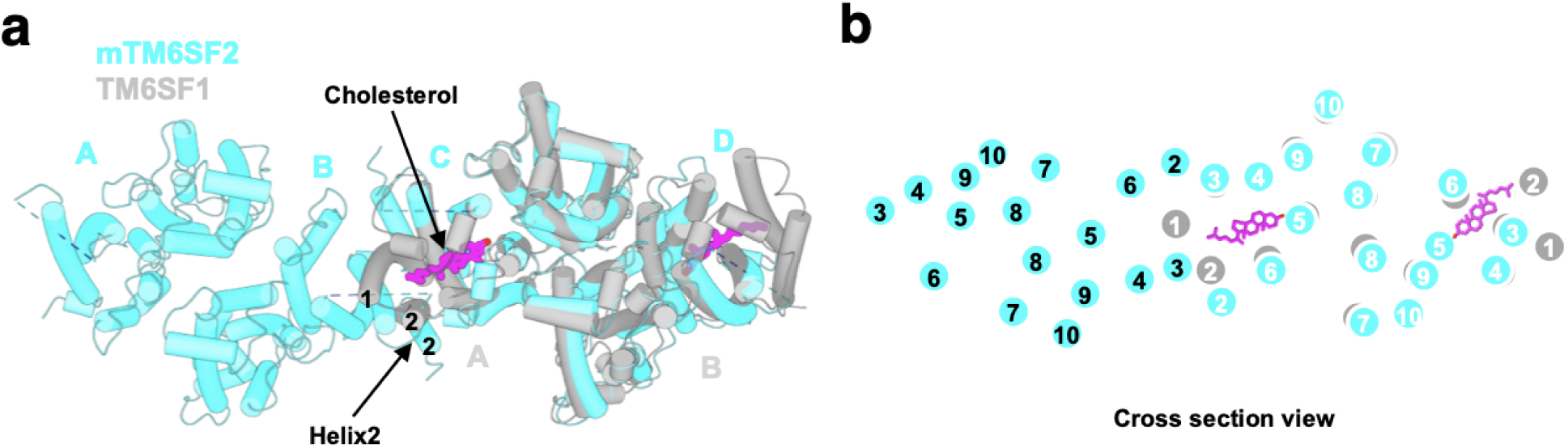
Structural alignment of mTM6SF2 tetramer and human TM6SF1. **a,** Superposition of the mTM6SF2 tetramer (cyan) with human TM6SF1 (grey; PDB: 10UP). Cholesterol bound to TM6SF1 is shown as a stick (magenta). **b,** Cross-sectional view of superposition showing the transmembrane helices and the location of cholesterol molecules.

**Extended Data Fig. 6.**
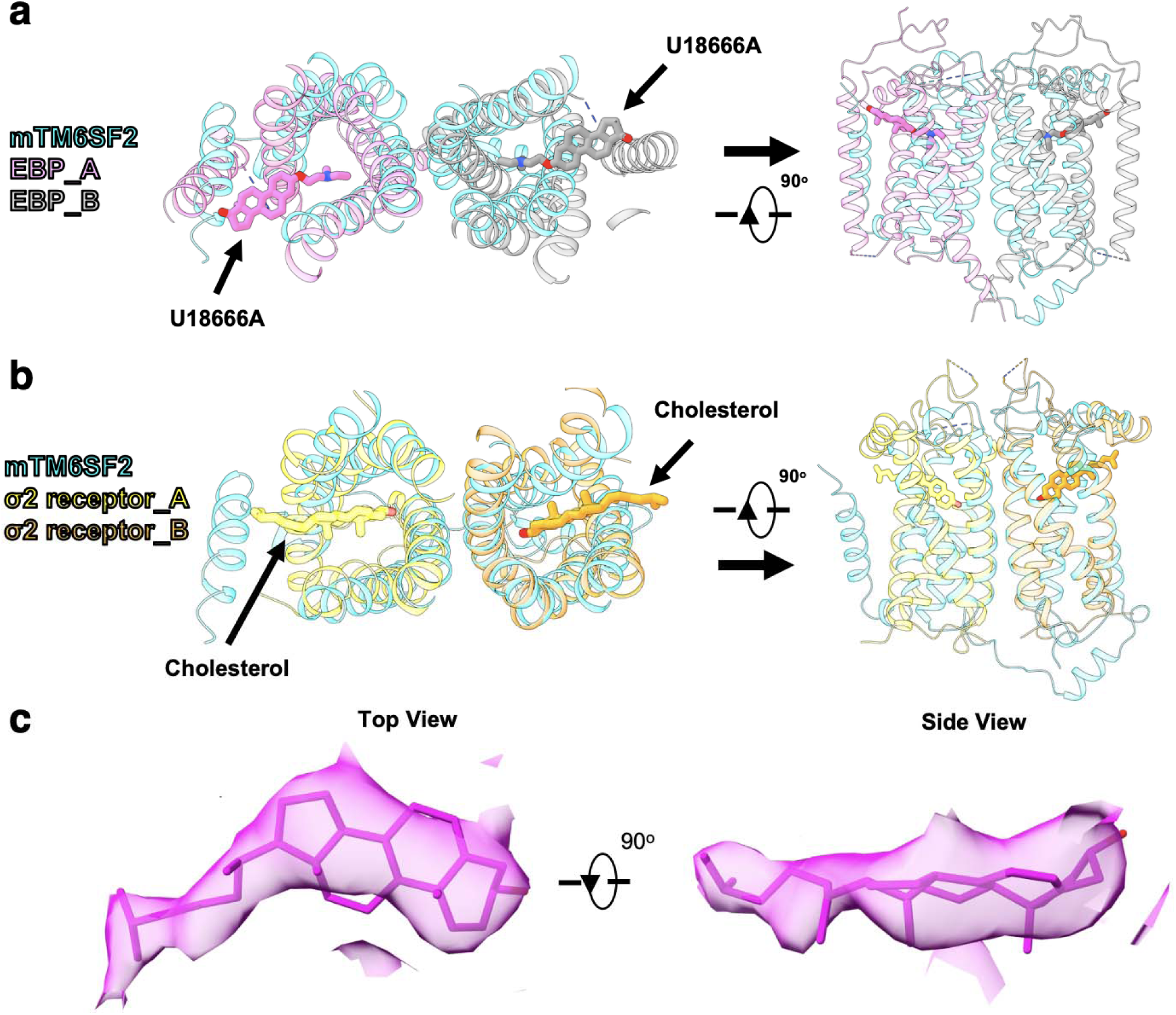
Structural comparison of mTM6SF2 with EBP and σ2 receptor. **a,** Superposition of mTM6SF2 (cyan) with emopamil-binding protein (EBP; PDB: 6OHT). EBP protomers A and B are shown in pink and grey, respectively. The sterol-like inhibitor U18666A bound to EBP is shown as a stick. Top (left) and side (right) views of the transmembrane helical bundle are shown. **b,** Structural alignment of mTM6SF2 (cyan) with the σ2 receptor (PDB: 7MFI). The σ2 receptor protomers A and B are shown in yellow and orange, respectively, with bound cholesterol molecules shown as sticks. Top and side views highlight the cholesterol-binding sites. **c,** Cryo-EM density corresponding to cholesterol.

**Extended Data Fig. 7.**
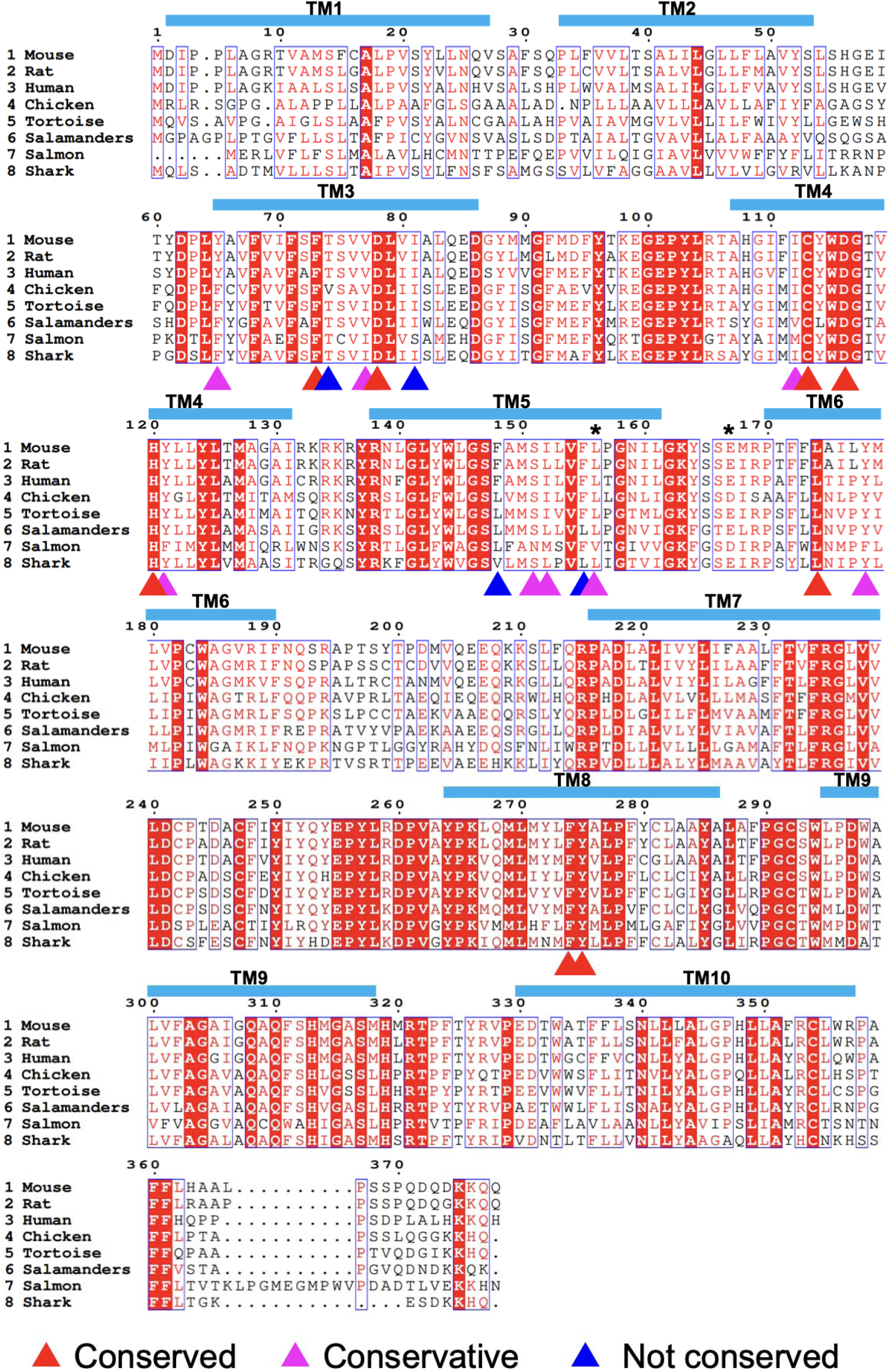
Sequence alignment of TM6SF2 homologs. Sequence alignment of TM6SF2 from mouse (NCBI Reference Sequence: NP_001280724.1), rat (NP_001121126.1), human (NP_001001524.2), chicken (NP_001376558.1), tortoise (XP_032655564.1), salamanders (XP_078526446.1), salmon (NP_001134092.1), and shark (UniProt: A0A4W3H7V8) is shown. Transmembrane helices are indicated above the aligned sequences. Conserved, conservatively substituted, and non-conserved residues are marked with red, magenta, and blue triangles, respectively. Residues L156 and E167 are marked with asterisks.

**Extended Data Fig. 8.**
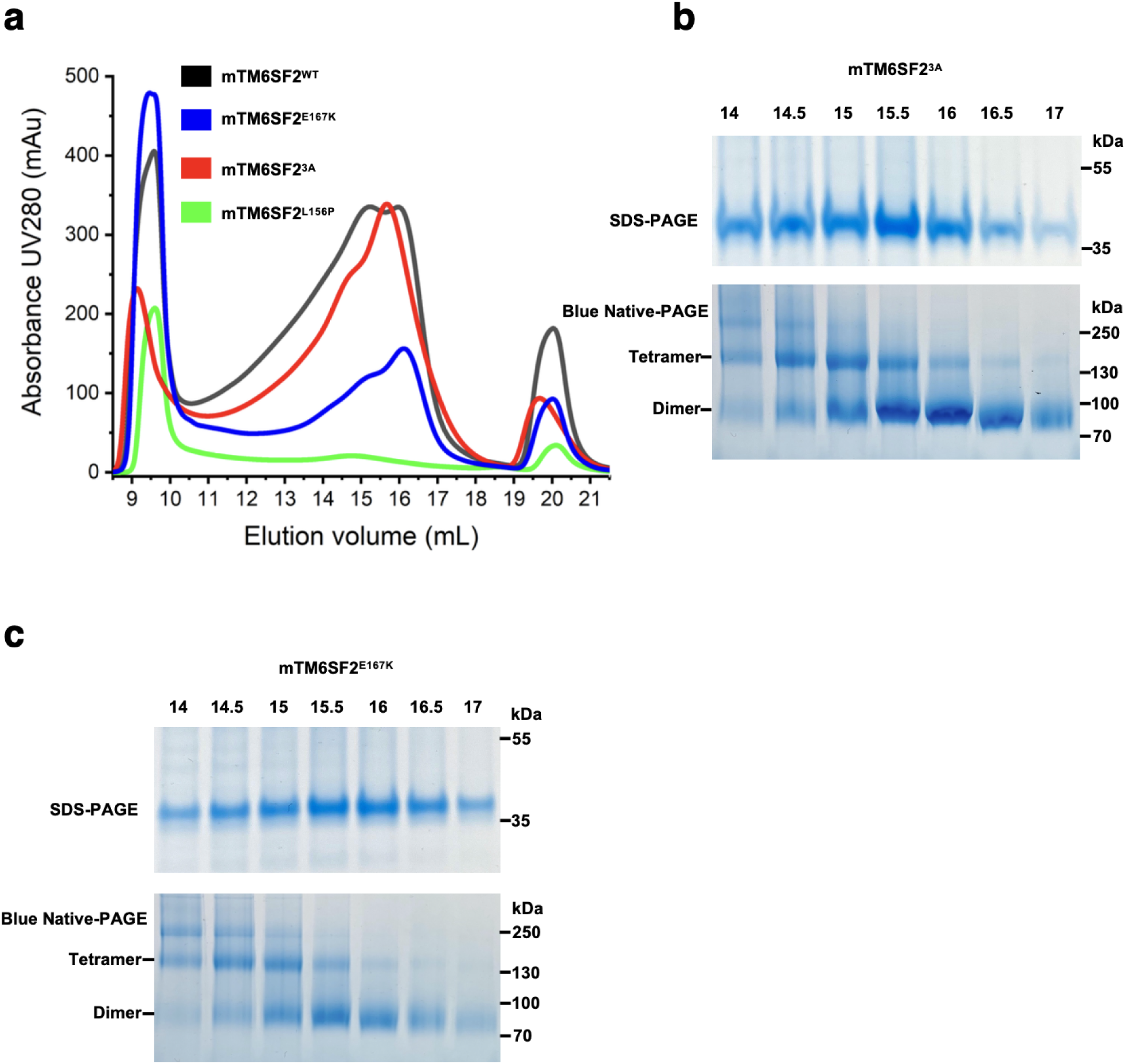
Expression and purification of the mTM6SF2 variants. **a,** Representative SEC profiles of mTM6SF2^WT^ (black), mTM6SF2^E167K^ (blue), mTM6SF2^3A^ (red), and mTM6SF2^L156P^ (green). **b, c,** SDS-PAGE and Blue Native-PAGE analysis of **(b)** mTM6SF2^3A^ and **(c)** mTM6SF2^E167K^.

**Extended Data Fig. 9.**
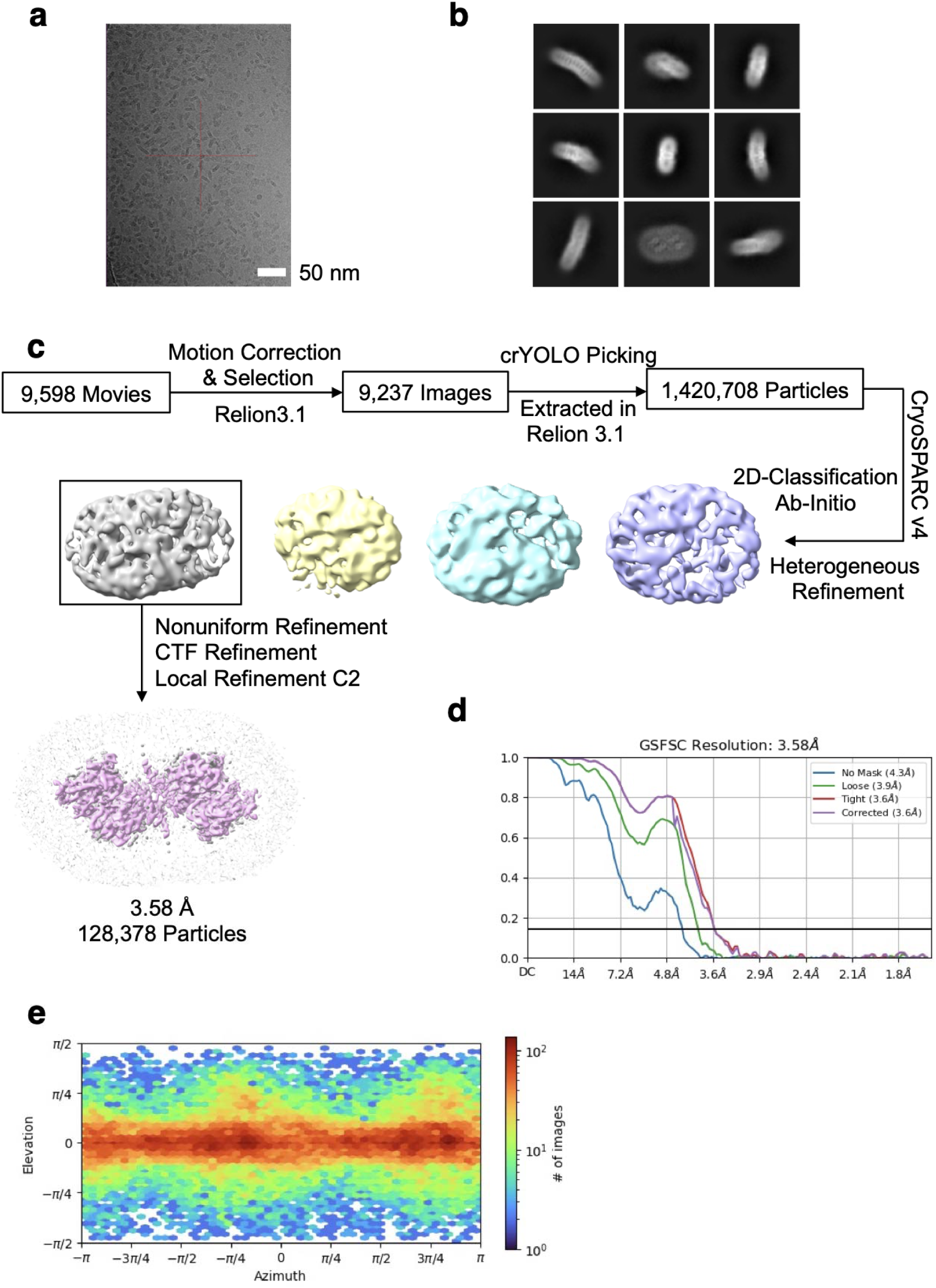
Cryo-EM analyses of tetrameric mTM6SF2^E167K^. **a,** Representative cryo-EM micrograph of tetrameric mTM6SF2^E167K^ in GDN micelles. Scale bar, 50 nm. **b,** Representative 2D class averages of tetramer with box size of 289.45 Å. **c,** Workflow of image processing for the mTM6SF2^E167K^ tetramer. **d,** Fourier shell correlation (FSC) curves between two half maps of the mTM6SF2^E167K^ tetramer. **e,** Angular distribution of particles used in the final 3D reconstruction of mTM6SF2^E167K^ tetramer.

**Extended Data Fig. 10.**
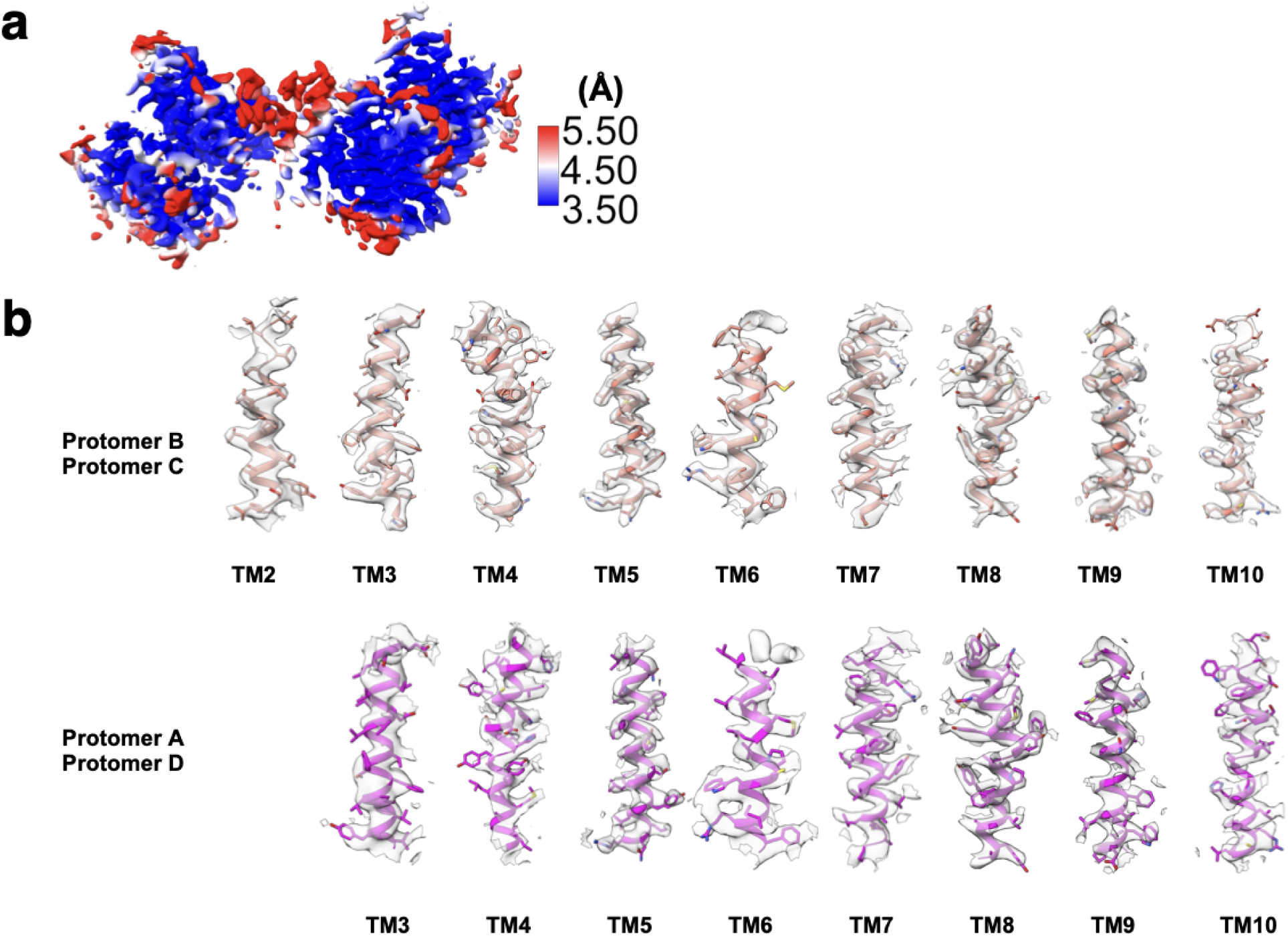
Cryo-EM density maps of tetrameric mTM6SF2^E167K^. **a,** Local resolution map of the mTM6SF2^E167K^ tetramer, colored according to local resolution estimated by cryoSPARC. The color gradient (blue-white-red) represents a resolution range from 3.5 Å to 5.5 Å. Cryo-EM density of the TM helices of **b,** protomer B and C and **c,** protomer A and D.

**Extended Data Fig. 11.**
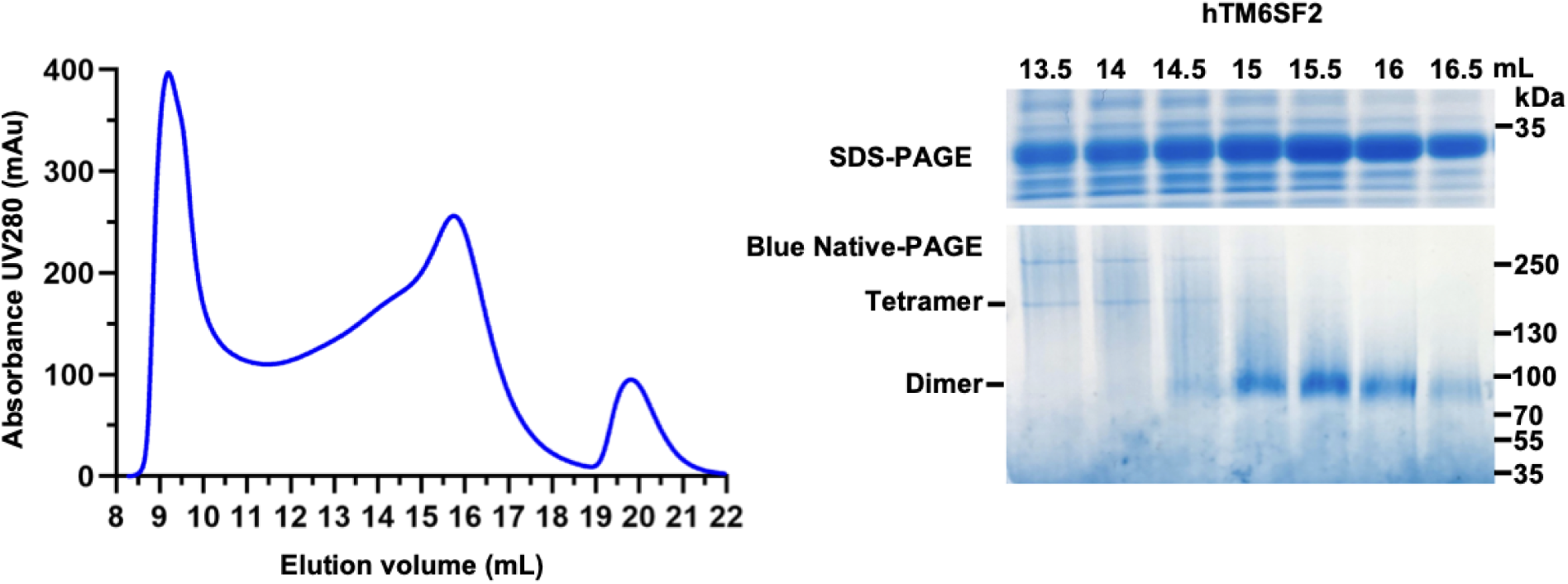
Expression and purification of human TM6SF2 (hTM6SF2). Representative SEC profile of purified hTM6SF2 using a Superose 6 Increase 10/300 GL column. SDS-PAGE and Blue Native-PAGE analysis of SEC fractions, with molecular weight markers indicated, are shown on the right

**Extended Data Table 1.**
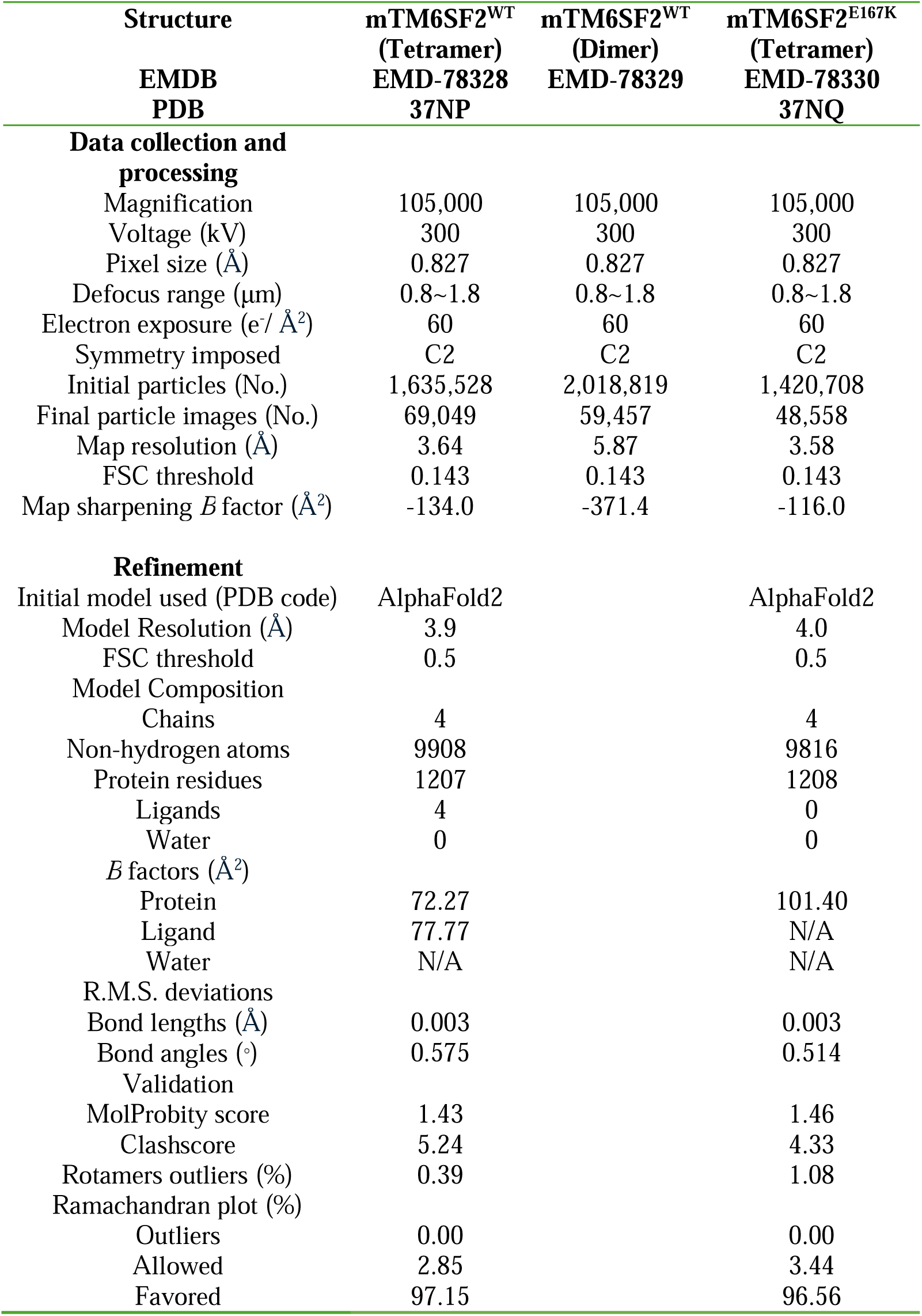
Cryo-EM data collection, refinement, and validation statistics.

