## Supplementary information for "TM6SF2 is a cholesterol-bound homomultimeric ER membrane protein that binds ApoB and is required for bulk lipidation of VLDL" (from another version of this preprint: version 1, 2026-08-18)

**Extended Data Fig. 1**

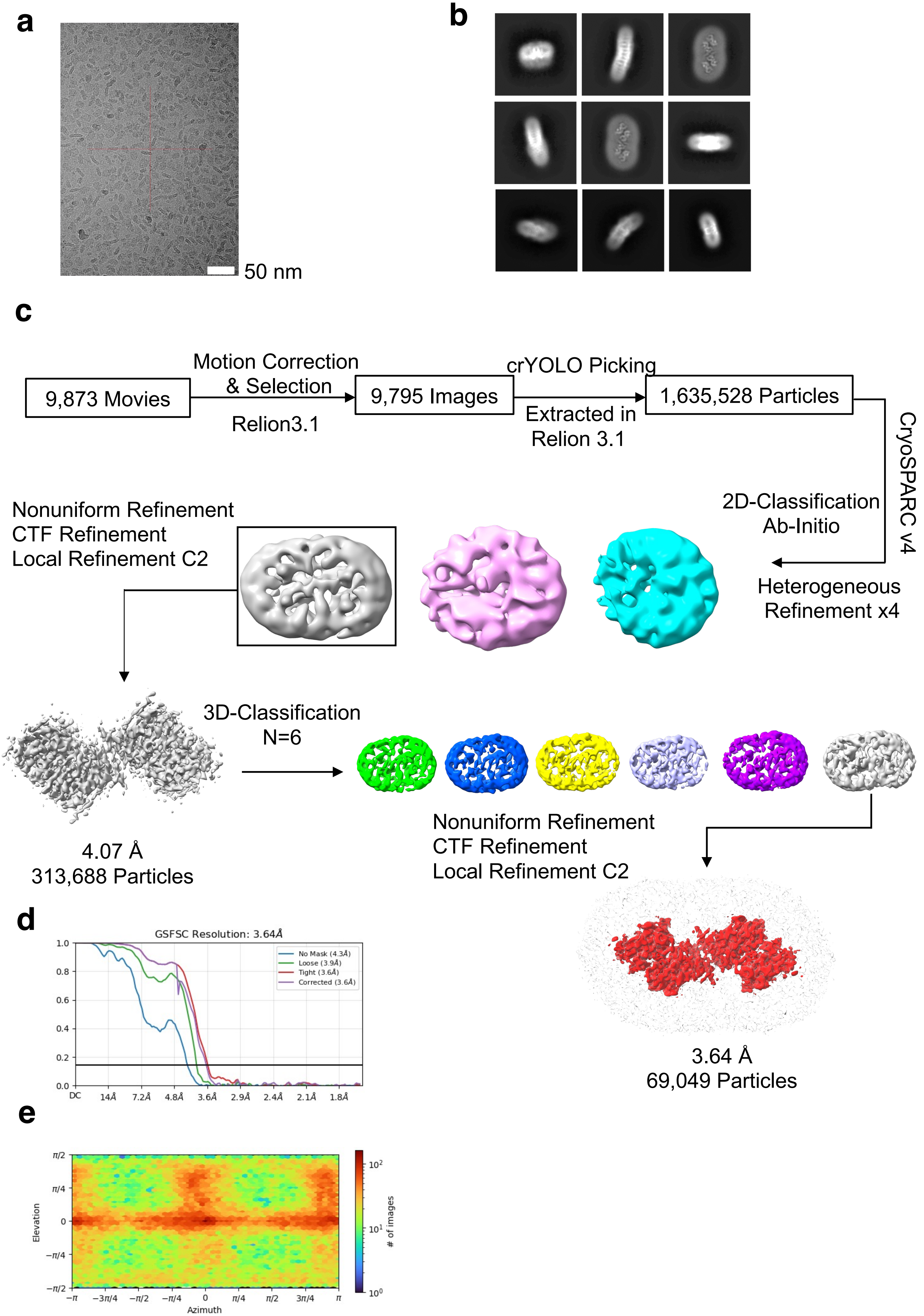

### Extended Data Fig. 2

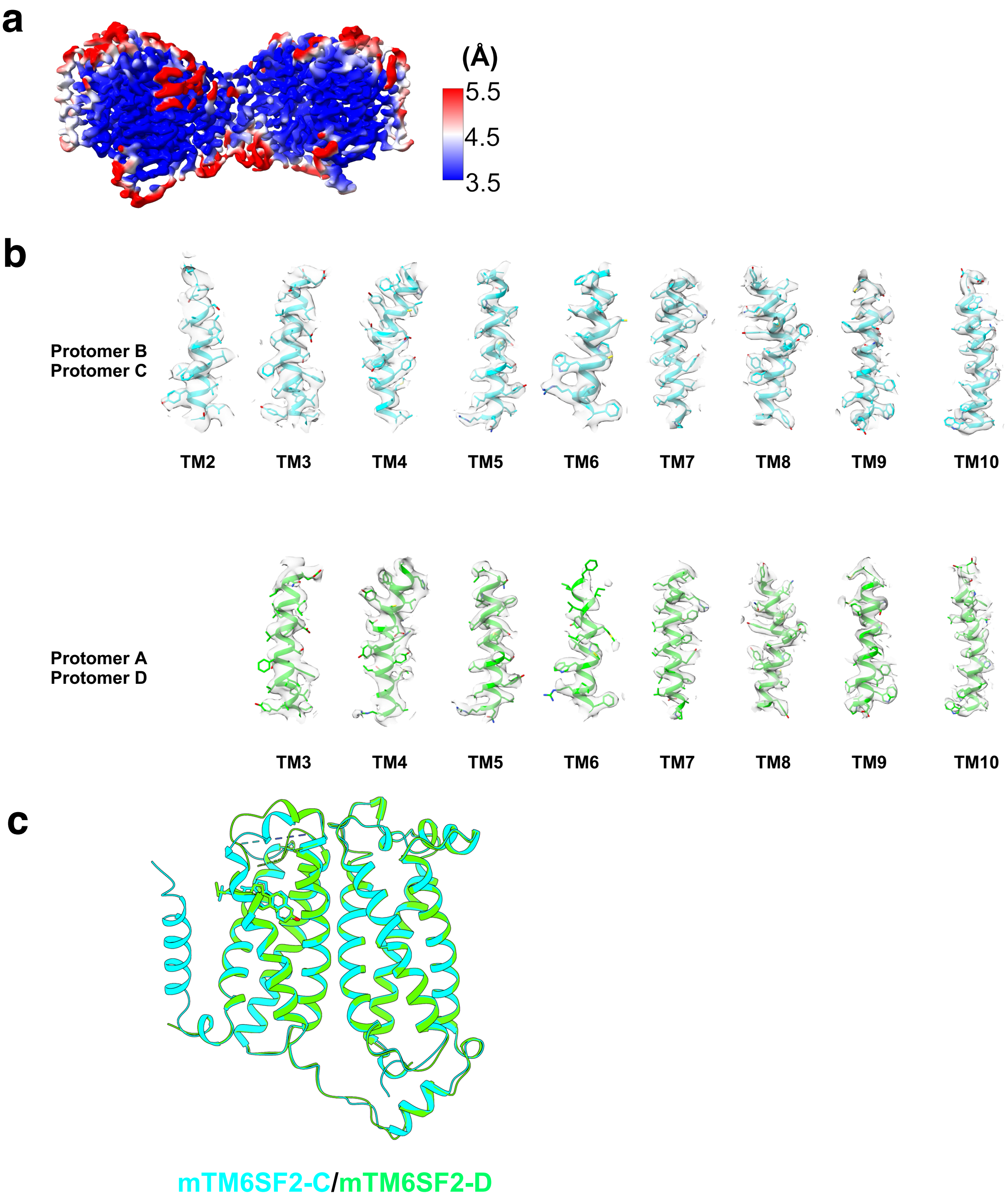

### Extended Data Fig. 3

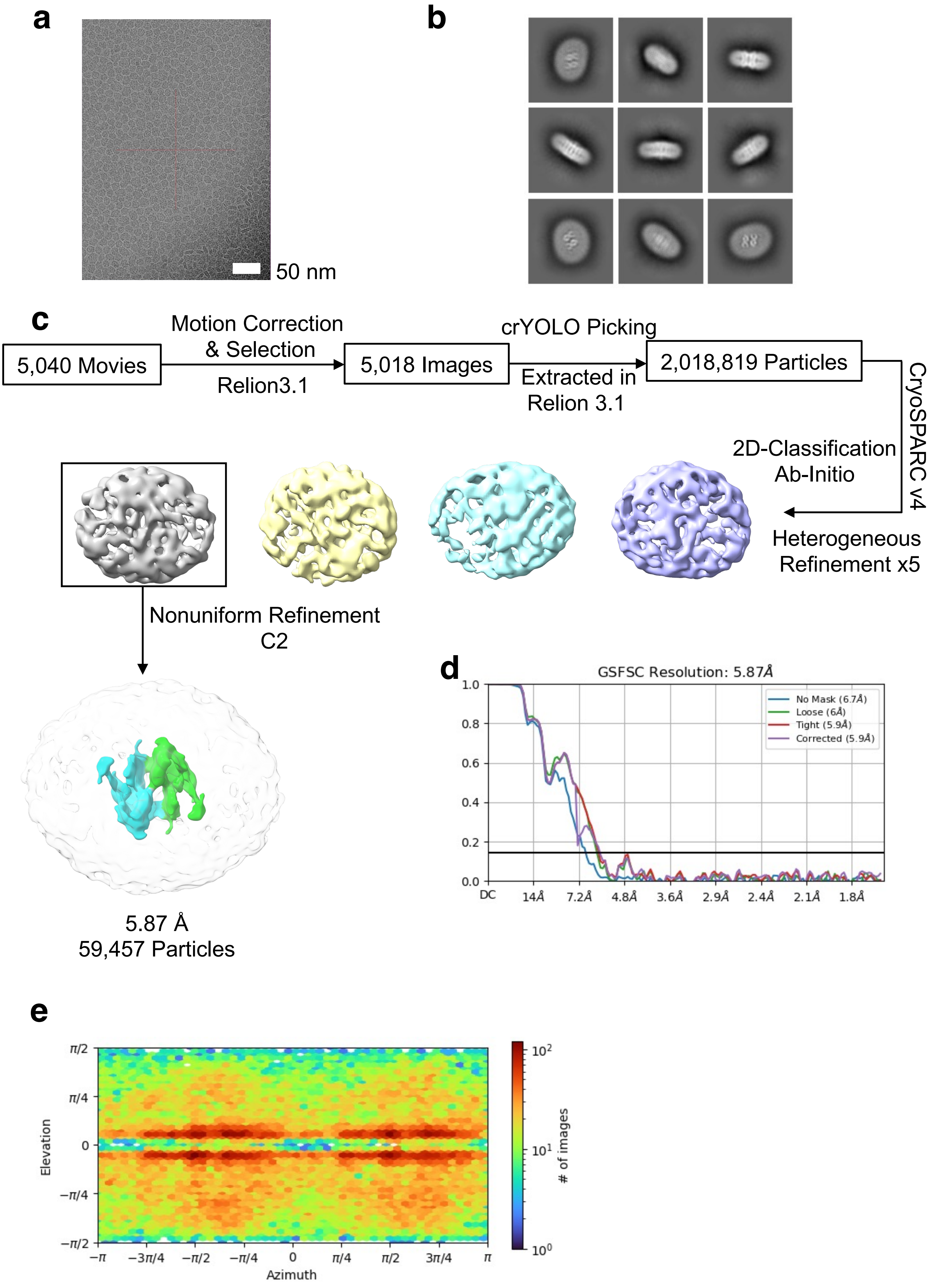

### Extended Data Fig. 4

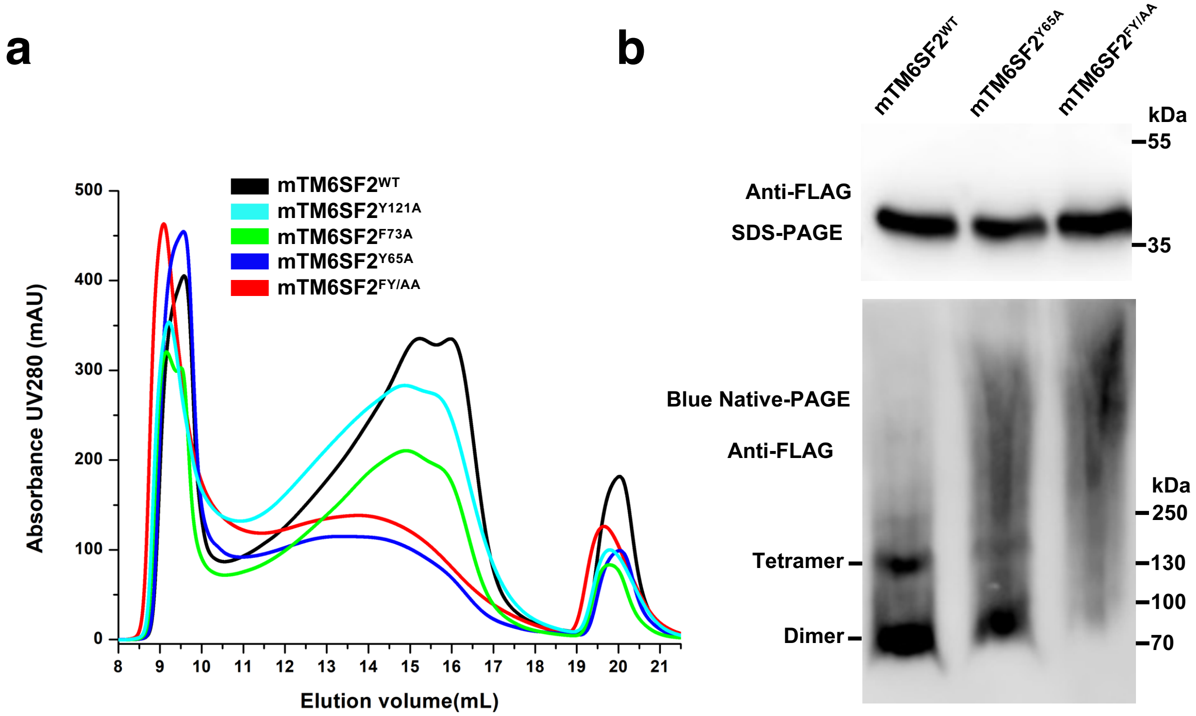

### Extended Data Fig. 5

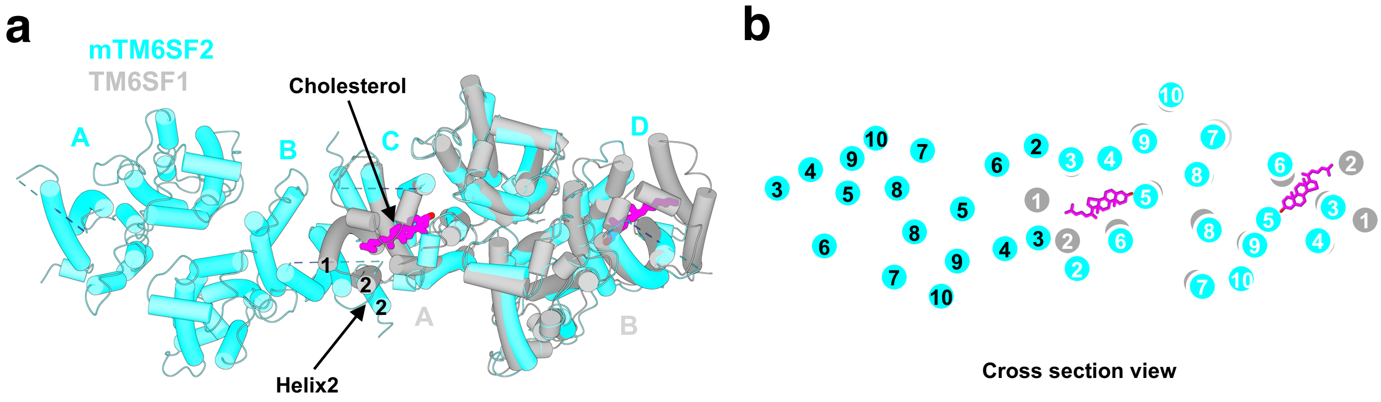

### Extended Data Fig. 6

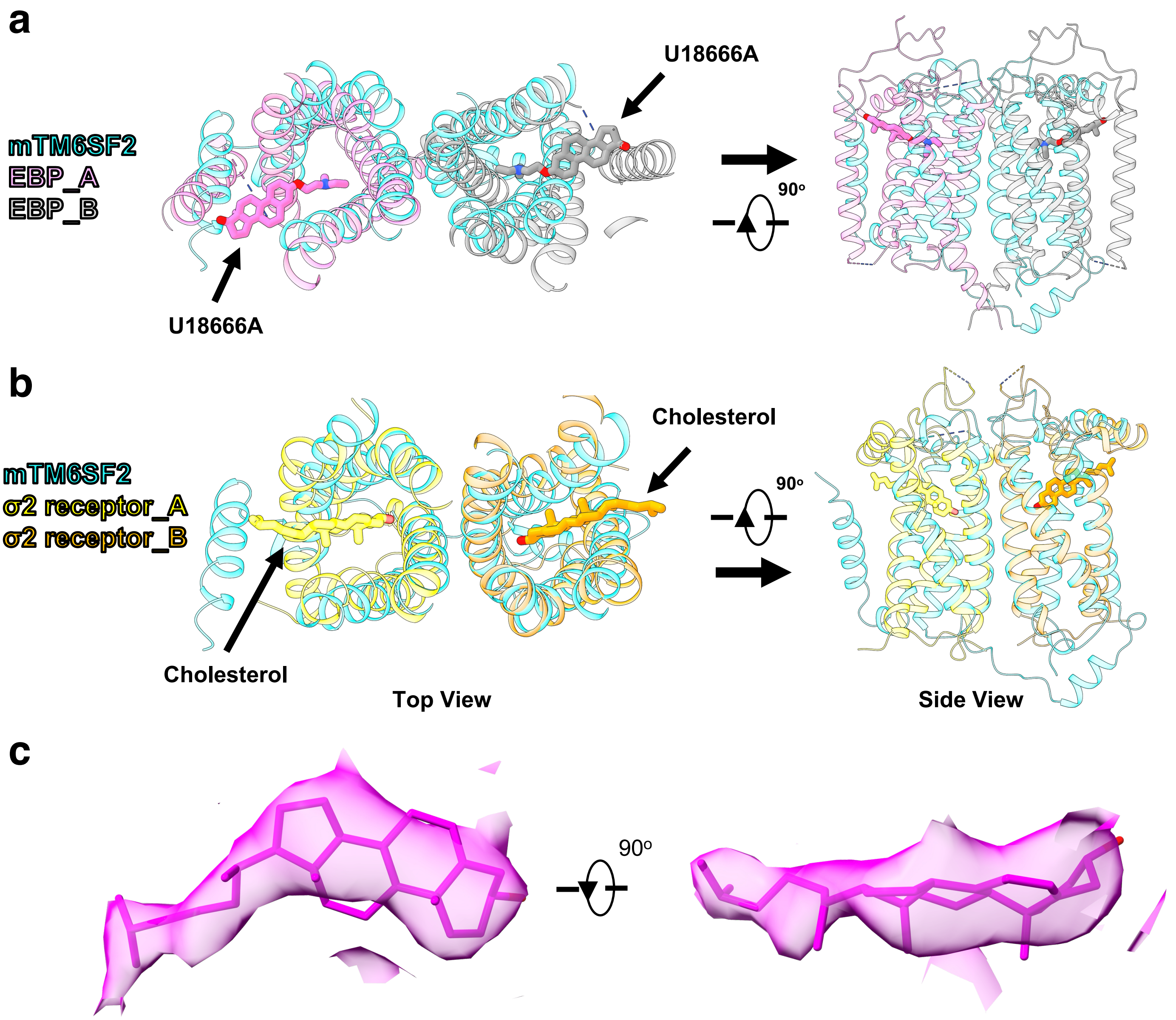

**Extended Data Fig. 7**

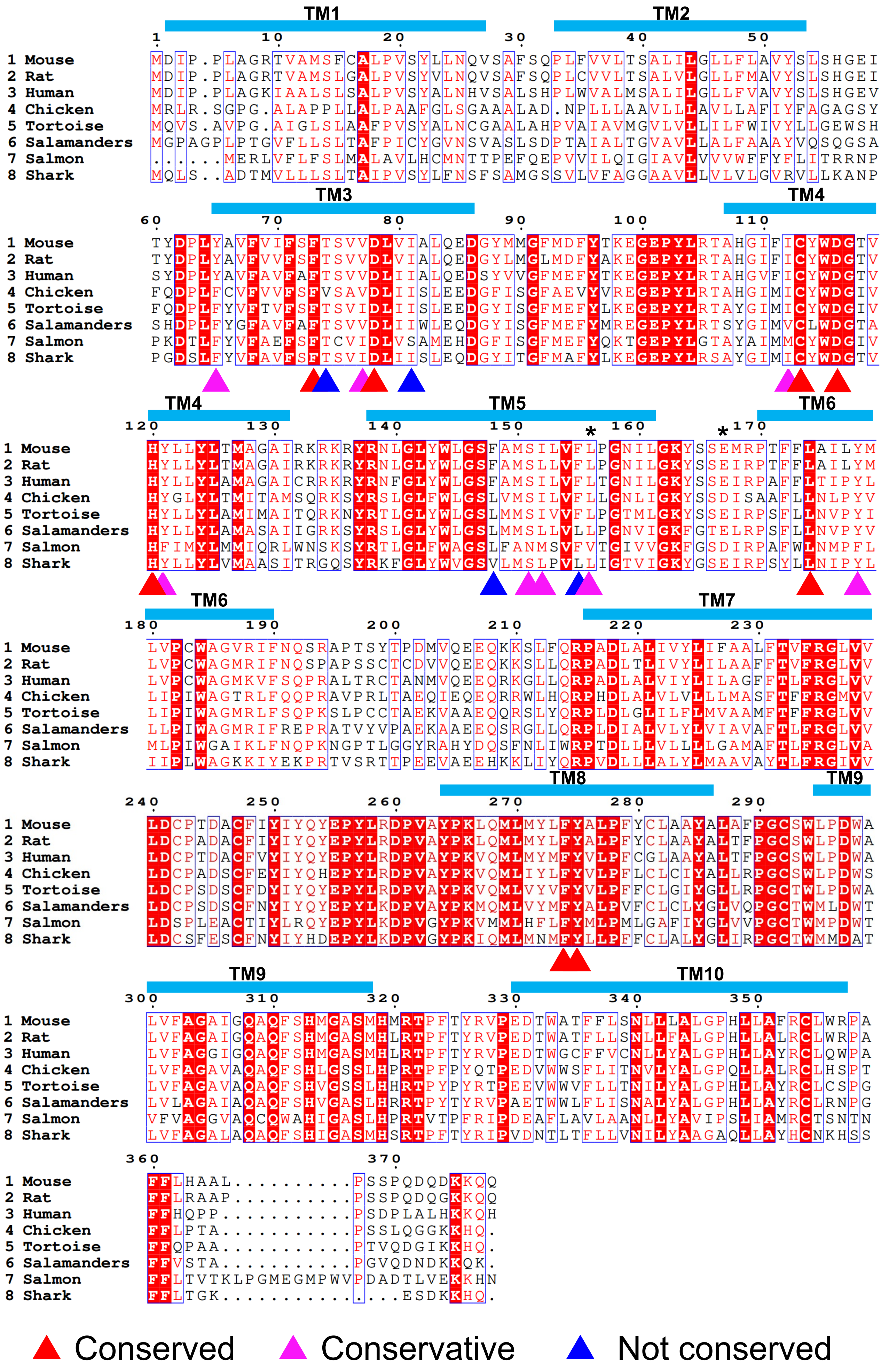

#### Extended Data Fig. 8

a

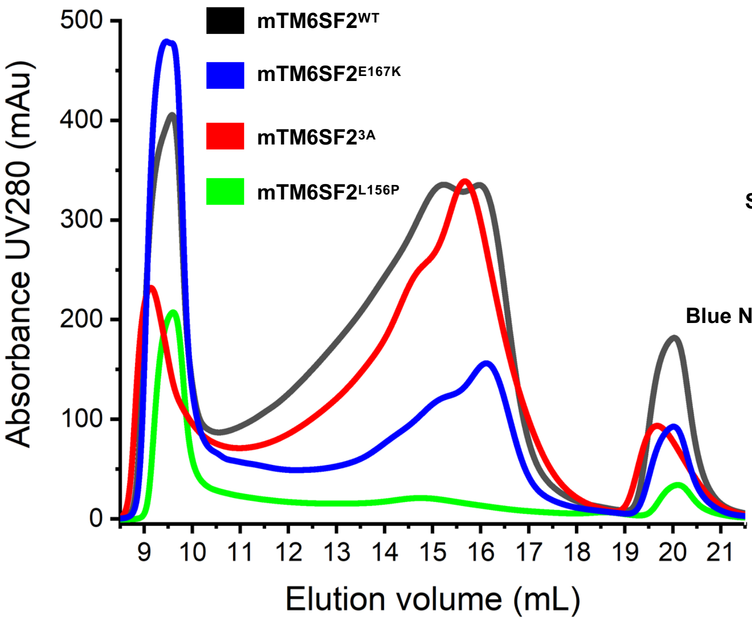

b

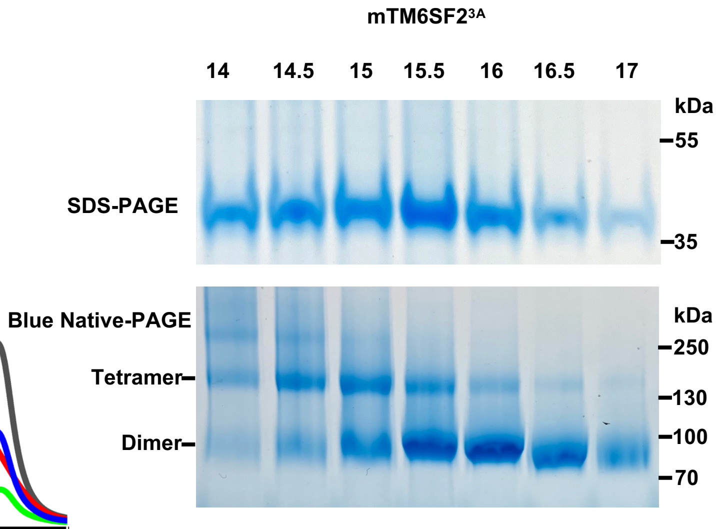

c

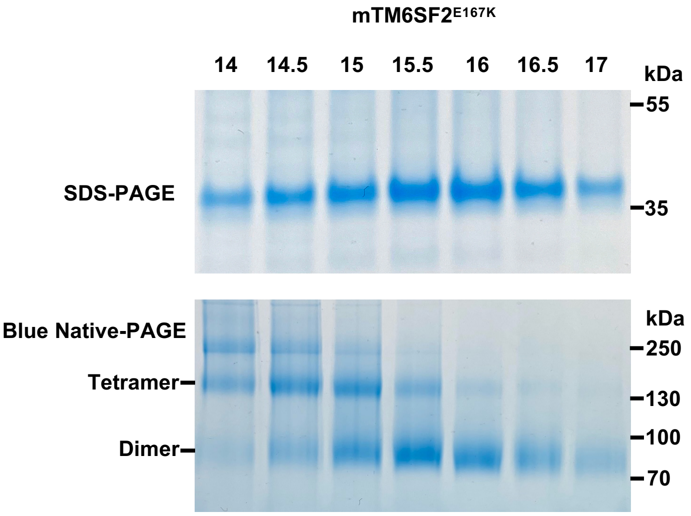

#### Extended Data Fig. 9

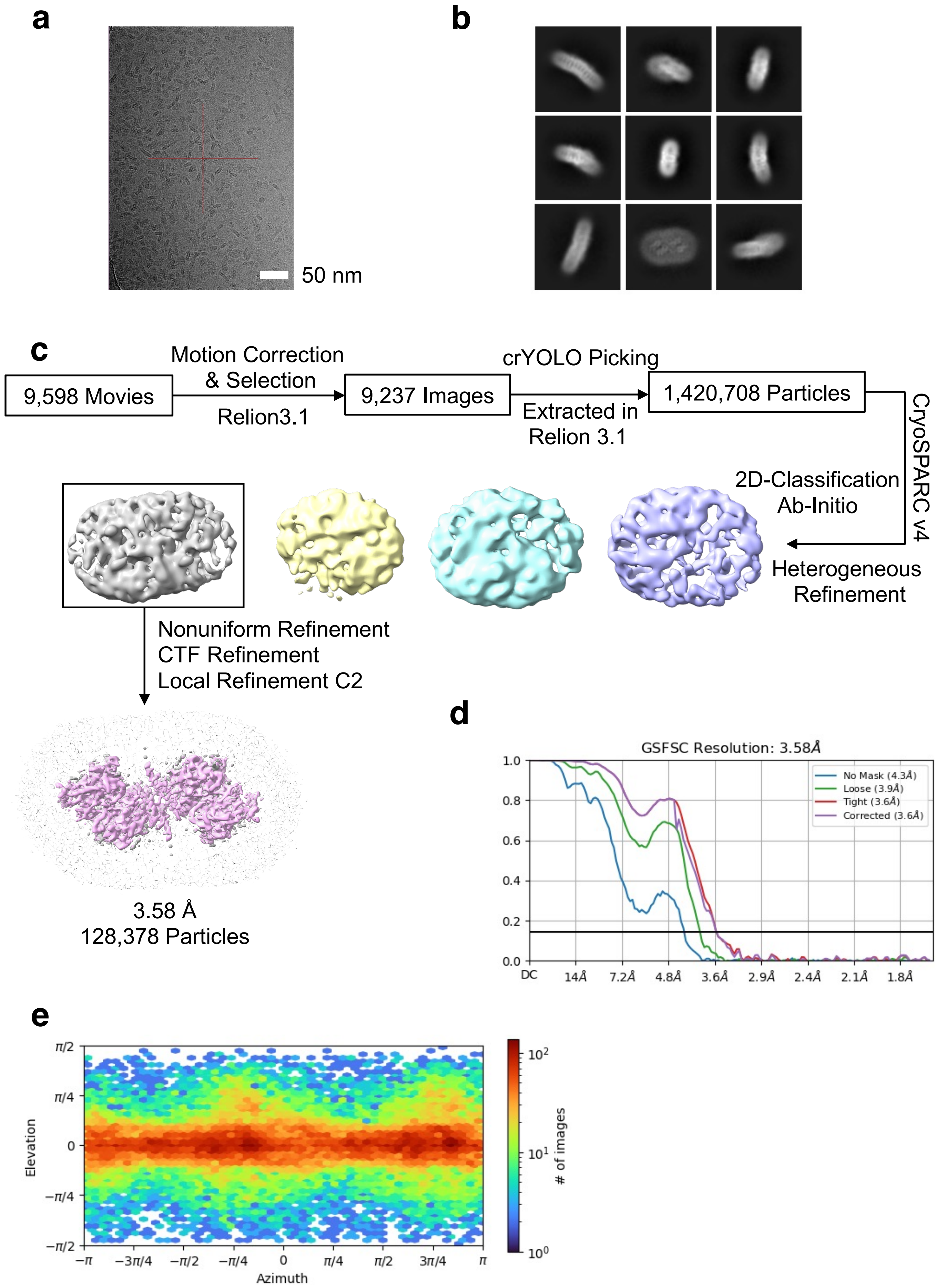

#### Extended Data Fig. 10

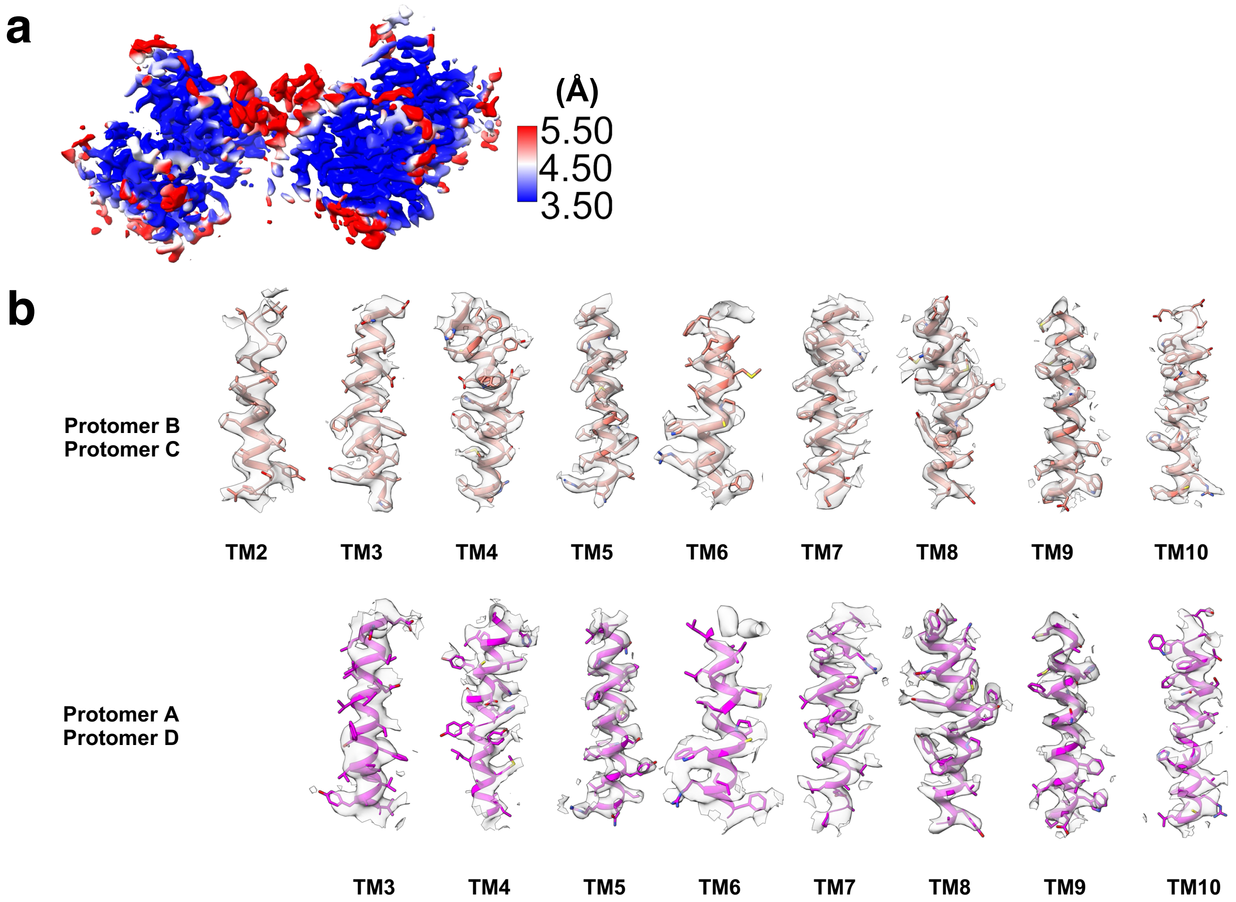

### Extended Data Fig. 11

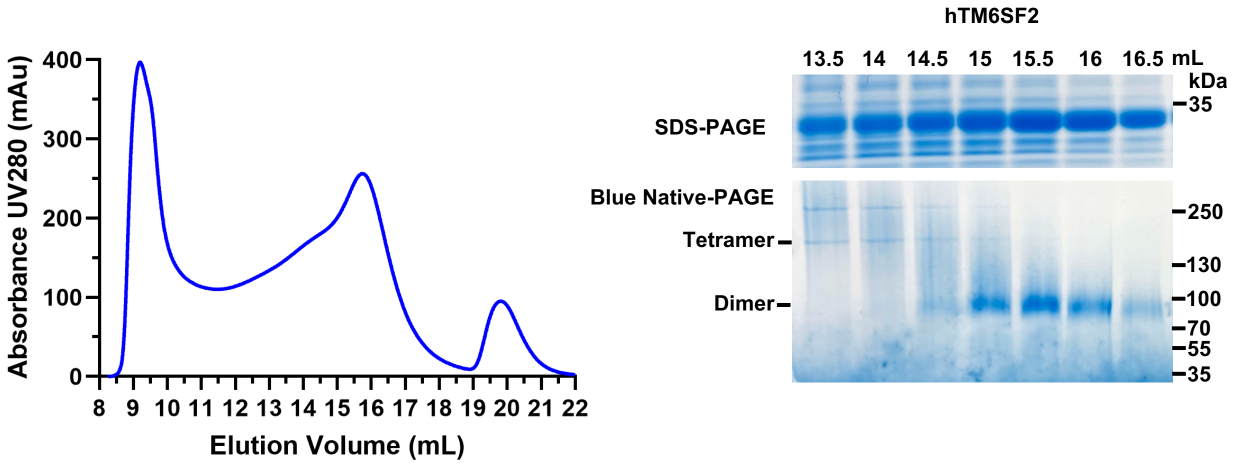

**Extended Data Table 1 Cryo-EM data collection, refinement, and validation statistics**

| **Structure** | **mTM6SF2$^{WT}$ (Tetramer)** | **mTM6SF2$^{WT}$ (Dimer)** | **mTM6SF2$^{E167K}$ (Tetramer)** |
|---|---|---|---|
| **EMDB** | **EMD-78328** | **EMD-78329** | **EMD-78330** |
| **PDB** | **37NP** | | **37NQ** |
| **Data collection and processing** | | | |
| Magnification | 105,000 | 105,000 | 105,000 |
| Voltage (kV) | 300 | 300 | 300 |
| Pixel size (Å) | 0.827 | 0.827 | 0.827 |
| Defocus range (μm) | 0.8~1.8 | 0.8~1.8 | 0.8~1.8 |
| Electron exposure ($e^-$/ Å$^2$) | 60 | 60 | 60 |
| Symmetry imposed | C2 | C2 | C2 |
| Initial particles (No.) | 1,635,528 | 2,018,819 | 1,420,708 |
| Final particle images (No.) | 69,049 | 59,457 | 48,558 |
| Map resolution (Å) | 3.64 | 5.87 | 3.58 |
| FSC threshold | 0.143 | 0.143 | 0.143 |
| Map sharpening *B* factor (Å$^2$) | -134.0 | -371.4 | -116.0 |
| **Refinement** | | | |
| Initial model used (PDB code) | AlphaFold2 | | AlphaFold2 |
| Model Resolution (Å) | 3.9 | | 4.0 |
| FSC threshold | 0.5 | | 0.5 |
| Model Composition | | | |
| Chains | 4 | | 4 |
| Non-hydrogen atoms | 9908 | | 9816 |
| Protein residues | 1207 | | 1208 |
| Ligands | 4 | | 0 |
| Water | 0 | | 0 |
| *B* factors (Å$^2$) | | | |
| Protein | 72.27 | | 101.40 |
| Ligand | 77.77 | | N/A |
| Water | N/A | | N/A |
| R.M.S. deviations | | | |
| Bond lengths (Å) | 0.003 | | 0.003 |
| Bond angles (°) | 0.575 | | 0.514 |
| Validation | | | |
| MolProbity score | 1.43 | | 1.46 |
| Clashscore | 5.24 | | 4.33 |
| Rotamers outliers (%) | 0.39 | | 1.08 |
| Ramachandran plot (%) | | | |
| Outliers | 0.00 | | 0.00 |
| Allowed | 2.85 | | 3.44 |
| Favored | 97.15 | | 96.56 |
